# Horizontally acquired HosA transcription factor bound with 4-hydroxy-benzoic acid exhibits unique tug-of-water dynamics

**DOI:** 10.1101/2024.04.24.590444

**Authors:** Arpita Goswami, Samee Ullah, José A. Brito

## Abstract

Pathogenic *Escherichia coli* (*E. coli*) causes serious illnesses in humans aided by several multiple antibiotic resistance regulator (MarR) transcription factors. Among these proteins, HosA provides support to these organisms by executing crucial roles in functions ranging from flagella motility to organic compound metabolism. The crystal structure of HosA from enteropathogenic *E. coli* is elucidated in this study along with conformational changes orchestrated by different amino acids due to para-hydroxy benzoic acid (PHB) binding in the hinge region. Structural analysis and extensive molecular dynamics simulation reveal role of a dynamic water molecule as a bridging entity in PHB bound structure which is not shown for any MarR structure yet. Also, it is shown that the HosA gene is transferred horizontally from *Shigella* to pathogenic *E. coli*, having 97.6% sequence similarity with an uncharacterized transcription factor from *Shigella dysenteriae Sd197*. This study may be promising to address several unanswered questions for the functioning of MarR transcription factors from infectious *E. coli* and design inhibitors to combat these pathogens.

## Introduction

Pathogenic *Escherichia coli* (*E. coli*) infections are a world-wide concern. They exhibit a broad spectrum of infections targeting host gastro-intestinal, urinary, and circulatory and nervous systems causing myriads of diseases. There are also several specific proteins in these pathogens which provide support for these infections. These are absent in non-infecting *E. coli* strains like MG1655, K12 and other laboratory strains like DH5α, BL21 etc. Transcription factor HosA is one such molecule present in pathogenic *E. coli*. There is report of these proteins playing major role in flagellar motility in enteropathogenic *E. coli O127:H6* which causes infantile diarrhea (1). The major structural protein flagellin (FliC) (2) which constitutes the core of flagellar motor is expressed less in the mutants devoid of HosA transcription factor (1). Both HosA and homolog SlyA come under MarR (multiple antibiotic resistance regulator) family of DNA binding proteins which serve wide range of functions (3). All these proteins are dimeric and have strikingly similar structure, most importantly the DNA binding domain as winged helix-turn-helix (3). The key recognition helix interacts with DNA by binding to major grooves while winged β-turn attaches to adjacent minor grooves. Several organic compounds (e.g. Salicylate, Ethidium bromide, 4-hydroxy-phenylacetate etc.) are known to disrupt this interaction by binding to a pocket above DNA binding region to cause local changes (3). This mechanism is universally common among all MarR family proteins from pathogenic *E. coli* and other bacteria like *Salmonella enterica, Klebsiella pneumoniae* etc (3).

Biochemical studies on HosA from uropathogenic *E. coli* UMN026 has shown hydrolyzed paraben, 4-hydroxy-benzoic acid (4-HBA) or para-hydroxy-benzoic acid (PHB) as a ligand for this protein (4). 4-HBA binding to HosA disrupts DNA binding (4). It was also shown that HosA in these bacteria are involved in auto repression as well as down-regulation of nonoxidative hydroxyarylic acid decarboxylase (HAD) operon involved in degrading organic disinfectants (4). One of the substrate of the biochemical pathway controlled by this operon is 4-HBA (4). It was proven biochemically that binding of 4-HBA to HosA significantly unregulated the operon by several folds which was otherwise switched off in untreated culture (4). Alongside, the palindromic DNA binding site of HosA was also identified which was downstream to transcriptional start site suggesting a steric blockade for movement of RNA polymerase (4). It was suggested that diffusion of hydrolysed paraben from extra cellular environment causes immediate capture by HosA bound to DNA to result dislocation and resumption of DNA transcription (4). Several studies utilized this mechanism to develop a bio-sensor device to detect PHB which is added as preservatives in commodities (5, 6).

All these studies underscores the importance to investigate the structure for HosA as apo-protein and as complex with 4-HBA. Since the sequences of HosA across pathogenic *Escherichia coli* are highly similar, any study on any one of them will unravel others also. We focus on HosA from enteropathogenic *E. coli O127:H6* which is known as a model organism to study these pathogens due to availability of whole genome sequence data (7). The analysis from Foldseek (8) and genome mapping indicates that this gene is acquired by enteropathogenic *E. coli O127:H6* from *Shigella dysenteriae Sd197*. Through structural perspective, we also exhibit a novel single water mediated tug-of-water dynamics playing major role in presence of 4-HBA.

## Materials and Methods

### Cloning and expression of HosA

*Escherichia coli* O127:H6 (strain E2348/69) HosA DNA sequence retrieved from NCBI (Reference genome: NZ_LT827011.1) was codon optimized for *Escherichia coli*. The sequence was further cloned in pET21a+ (Novagen) between NdeI and XhoI restriction sites. The synthesized gene had a 6X-His tag from the vector at C-terminus before the stop codon. Further, the cloned plasmid was transformed into BL21(DE3) cells (Novagen). Primary culture in LB (Luria Bertani) broth (HiMedia) from the transformed colony was grown overnight at 37°C and 200 rpm. 5 ml of secondary LB culture from the same (1 % inoculum) was grown at 37°C until OD reached 0.6. 1 mM IPTG was added thereafter and further the culture was grown for 4 hours at 37°C. In parallel, 5 ml uninduced culture was maintained using the same conditions but without IPTG. The overgrown cultures were pelleted, resuspended in a lysis buffer (50 mM Tris pH 8, 150 mM NaCl, 100 μM PMSF) and lysed by sonication. Post sonication lysates were centrifuged at 12,000 rpm at 4°C and 12% SDS-PAGE gel was run to observe solubility profiles of the protein after expression.

### Overexpression and Purification of HosA

The overexpression of the culture was done in 2 liter LB broth using optimized expression conditions as described above. The supernatant from lysed & centrifuged cell pellets was loaded in the Ni-NTA column (HisTrap FF, Cytiva). The washing and purification was done as per manufacturer’s protocol. The eluted protein in one fraction was >99% pure. This fraction was concentrated to 500 µl and used for crystallization. The final concentration of the protein as estimated from the BCA test (Sigma Aldrich) was 15.4 mg/ml.

### Crystallization of HosA

The apo-protein was crystallized in microbatch under oil method using 96 well vapor batch plates (Douglas) as 1:1 ratio (protein:precipitant). The usual screens from Hampton, Rigaku and Jena were used. Best quality crystals were obtained in 4°C and low pH conditions with ammonium acetate and PEG3350. The conditions were further optimized by grid setup. Final diffraction quality crystals were grown in 160 mM ammonium acetate, 100 mM Bis-Tris (pH 5.6), 27% PEG3350 (w/v). The ligand bound crystals were also grown in a similar method at 4°C. 4-hydroxy benzoic acid (SRL) was added to the protein *in situ* as 1:2 (protein:ligand) ratio. Optimum crystals were grown after adding calcium chloride in ammonium acetate based conditions. Final 4-HBA bound crystals were in 180 mM ammonium acetate, 12 mM calcium chloride, 50 mM sodium cacodylate (pH 6.5), 10% (w/v) PEG 4000.

### Diffraction and data collection

The apo and 4-HBA bound crystals were scooped in cryoloop (Hampton) with 25% ethylene glycol as cryoprotectant added to the respective mother liquors. After flash freezing in liquid nitrogen, the crystals were subjected to diffraction in Rigaku Micromax-007 diffractometer (X-ray facility, CSIR-CCMB, India) with MAR 345 image plate detector and rotating anode X-ray (CuKα) generator set at 50 kV, 100 mA voltage. Preliminary spacegroup analysis in *iMOSFLM* (9) from the first few frames collected showed the space group as P4 for both Apo and ligand bound crystals. Accordingly, the data collection tasks were set up to get >99% completeness in ∼ 5 hours. In parallel, diffraction was also carried out in INDUS synchrotron (RRCAT, Indore, India). For this, a wedge of 117° of data was collected from a single crystal at a wavelength of 0.979 Å and an oscillation angle of 1° per image.

### Structure solving

Indexing at first pointed to the P4_2_11 group by POINTLESS (*iMOSFLM*) (10,11) followed by scaling in *AIMLESS & CTRUNCATE* (*iMOSFLM*) (11,12) without any crystal defects (Twinning, Pseudo-translation etc.). Since, the presence of screw axis led systematic absences are known in this group. Further correction was obtained while doing molecular replacement in *PHASER* (13) in CCP4i suite (14, 15) by selecting analysis of all possible groups in P4. This conveniently confirmed the space group as P4_3_22. The template in *PHASER* was the C chain from the hexameric model of 6PCP (*Bordetella bronchiseptica* BpsR with 28% identity to HosA) (16). The single solution obtained from *PHASER* was subjected to model building in *AUTOBUILD* (*PHENIX*) (17) using multiple non-crystallographic symmetry (ncs) options (= 1, 2, 3). A well-built model was obtained only in ncs=3 after a single cycle of model building. Final model for refinement was obtained after multiple cycles of model building from this initial model with ncs =1. Further, the raw reflection data was reindexed (*REINDEX, CCP4i*) (14, 15), scaled (*SCALA, CCP4i*) (14, 15) and this data was used in refinement (*phenix*.*refine* (18) to prevent model bias. The accuracy of the reindexed intensities were further verified by cross-checking in *phenix*.*xtriage* (19) which otherwise showed anomalies in original indexed data. The refinement was carried out until there were no structural defects and good refinement values. All the structures solved by this methodology are submitted to RCSB Protein Data Bank (RCSB PDB) (http://wwpdb.org/) (20) with following ids: 8PQ4, 8AGA, 8WSV, 8XB7 and 8XZU. In parallel, structure solving was also carried out using *autoPROC*/*STARANISO* pipeline (Global Phasing) (21, 22) for comparative analysis.

### autoPROC/Staraniso pipeline analysis

Structure solving was carried out using autoPROC/*STARANISO* pipeline on data collected for PHB bound HosA crystal from Indus synchrotron for comparative analysis. Diffraction data were indexed, integrated and isotropically or anisotropically scaled with *XDS* (23) and *AIMLESS* (12), or *XDS* (23) and *STARANISO* (22), respectively, both within the *autoPROC* data processing pipeline (21). Space group was assessed with *POINTLESS* (10) from the *CCP4* suite of programs (14,15), and data quality was assessed with the *phenix*.*xtriage* (19) tool within the *PHENIX* suite of programs (24). The structure solved by molecular replacement (MR) with *PHASER* (13) as implemented in *PHENIX*, using one chain (monomer) of the X-ray structure of the HosA transcriptional regulator from enteropathogenic *Escherichia coli* O127:H6 (strain E2348/69) (PDB 8PQ4) as the search model, devoid of any cofactors, solvent molecules, and other ligands. Iterative model building and refinement were carried out in a cyclic manner with *phenix*.*refine* (18), *BUSTER-TNT* (25) and *COOT* (26), until a complete model was built and refinement convergence achieved. The final model was validated with *MolProbity* (27) within *PHENIX* with the atomic coordinates and structure factors deposited in the Protein Data Bank under accession code 8YCV. Final data processing (both isotropic and anisotropic statistics are reported), and refinement statistics are depicted in Supplementary Table 1.

### Computational biology analysis

Foldseek search was carried out on *Escherichia coli* O127:H6 (strain E2348/69) HosA protein sequence using default parameters. The top 18 hits were selected. The genome comparisons were done in Snapgene software (https://www.snapgene.com/). PDBePISA (Proteins, Interfaces, Structures and Assemblies) (28) and APBS (Adaptive Poisson-Boltzmann Solver) (29) analyses were carried out from respective servers.

### Molecular dynamics methodology

Detailed in Supplementary document.

## Results

### HosA apo-protein structure is similar with other MarR proteins

The alignments across *Escherichia coli* HosA showed high sequence conservation (>70% identity) among the proteins while structural alignments with *Escherichia coli* MarR exhibited ∼100% structural identity (Supplementary Fig. 1 & 2). The monomer structure has all a-helices and winged β-sheet formed by antiparallel strands as observed across MarR family transcription factors. Furthermore, a distinct local kink was observed in HosA winged β-sheet (Supplementary Fig. 2). PDBePISA analysis showed the molecule with 2 subunits (Supplementary Fig. 3). These two subunits correlate with each other by a 2-fold crystallographic rotation axis (Supplementary Fig. 3A & 3B). The solvent accessible surface area for the monomers was 13810 Å^2^ which got reduced to 4761.2 Å^2^ upon dimer formation. The solvation free energy gain upon formation of the assembly (ΔG^int^) was-31.1 kcal/mole indicating energy favorable association. While the free energy of assembly dissociation (ΔG^diss^) and the rigid-body entropy change at dissociation (TΔS^diss^) was 29.7 kcal/mol and 12.3 kcal/mol respectively indicating thermodynamic stability. Therefore, a stable crystallographic dimer upon close association of oppositely charged surfaces from two molecules was possible (Supplementary Figure 3C). This was noted in other MarR proteins also (30).

### HosA is horizontally acquired by EPEC from Shigella *dysenteriae Sd197*

It is widely known that the MarR family of proteins have high structural similarity but very low sequence similarity. We also observed the same in conventional structural alignments of experimental protein structures from PDB with HosA which showed myriads of MarR proteins from diverse bacteria with high structural similarity but little sequence similarity (data not shown). But when a Foldseek search was carried out there was 97.6% sequence identity with putative transcription factor from *Shigella dysenteriae Sd197* (Fig.1A). In fact, there was remarkable sequence identity (>70%) with homologous putative transcription factors from three enterobacteria (*Shigella, Salmonella* and *Klebsiella*). The striking finding from Foldseek can be attributed to AlphaFold coupled search feature which enabled structural information of hundreds of uncharacterized proteins to be available for Foldseek platform. The high identity of EPEC HosA with homologous protein from *Shigella dysenteriae Sd197* indicates that this was acquired by horizontal transfer from *Shigella* to EPEC. Further genome analysis in Snapgene software confirmed this finding by mapping flanking genes surrounding HosA (EPEC) (Fig.1B) and HosA like transcription factor (*Shigella*) (Fig.1C) as highly identical and reversely oriented indicating homologous recombination. One of the flanking gene (RNA polymerase sigma factor; RpoS) was more identical to homologous gene of *Salmonella enterica* pointing towards multiple horizontal transfer events. The significant cross-bacterial identity of hosA gene, surrounding locus and the high conservation profile of HosA among different pathogenic *E. coli* indicates the essentiality of the same in pathogenic γ-proteobacteria for surviving in the same environmental niche as cohabited by enterobacterial pathogens. Similar property was also displayed by homolog *E. coli* SlyA, with a 91% sequence identity to SlyA in *Salmonella Typhimurium* (31).

**Fig. 1.**
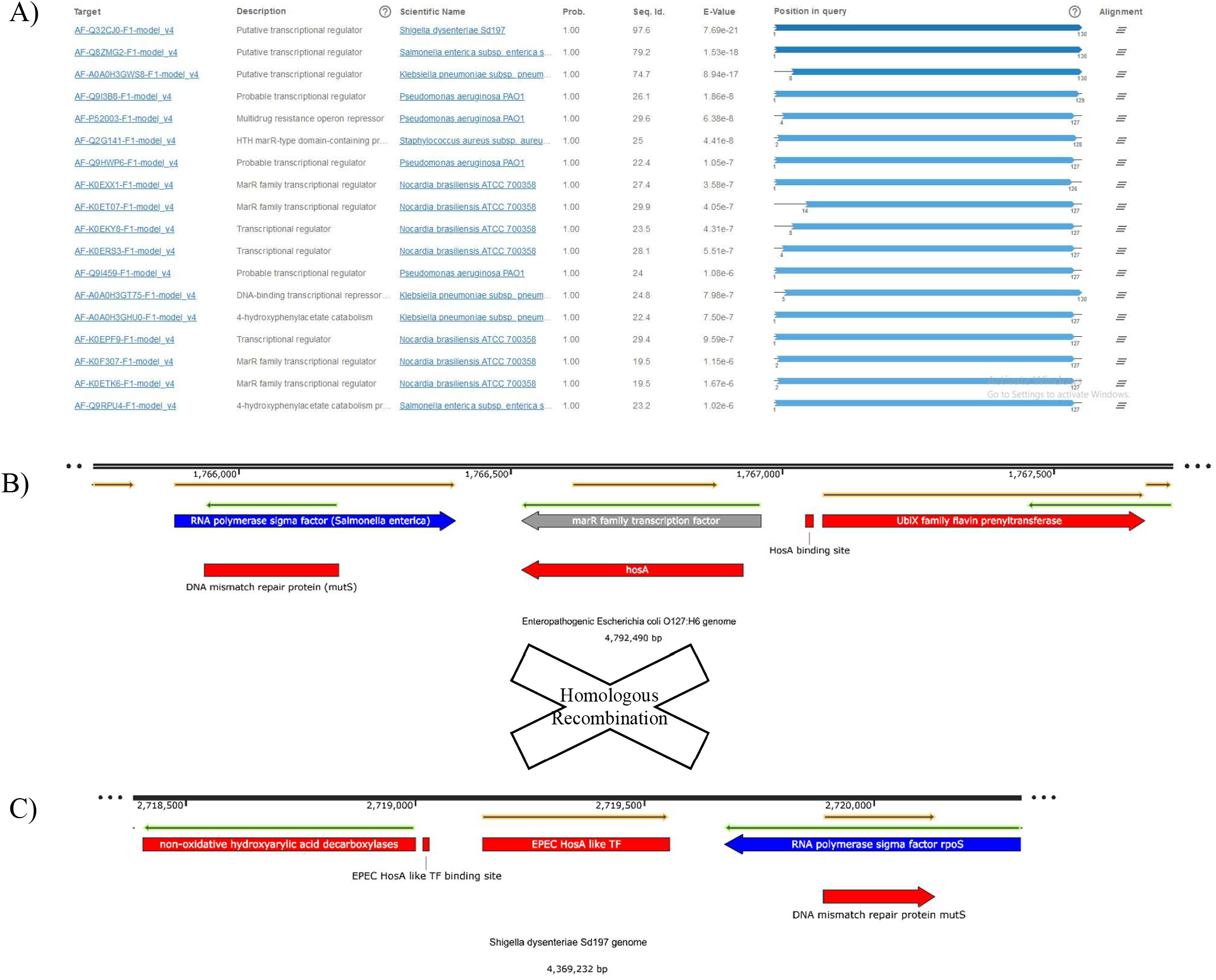
HosA is acquired by EPEC via horizontal transfer from Shigella dysenteriae Sd197. A) Foldseek analysis reveals a remarkable identity (97.6%, Top hit) between EPEC HosA protein sequence and a putative transcription regulator from *Shigella dysenteriae* Sd197, facilitated by AlphaFold structure (AF-Q32CJ0-F1-model_v4) as embedded in Foldseek algorithm. Additionally, significant identity (>70%) is observed between HosA and two other enterobacterial (*Salmonella enterica* and *Klebsiella pneumoniae*) homologs. B) Location of HosA gene (hosA, classified under *E. coli* marR family transcription factors) in EPEC genome aligns with C) EPEC like HosA transcription factor (TF) locus in *Shigella dysenteriae* Sd197 genome. The genes located on either sides of HosA are highly identical between the two genomes. The genes including hosA which are strikingly identical between EPEC and *Shigella* are coloured in red, while RNA polymerase sigma factor (rpoS) protein sequence is found slightly more identical with corresponding RpoS sequence from *Salmonella enterica*, hence coloured as blue. This suggests a simultaneous acquisition of genes by EPEC from *Shigella* and *Salmonella* by horizontal transfer. The orientations of the flanking genes of hosA are inverted between EPEC and Shigella genomes, indicating at least two homologous recombination events between EPEC and both *Shigella* and *Salmonella*.

### The binding of 4-hydroxy benzoic acid ligand happens by Aspartate110 from one subunit but requires presence of other subunit for retention

In order to explore the mechanism of 4 hydroxy benzoic acid (PHB) binding to HosA protein and downstream cascades, soaking of apo-protein crystals with PHB was carried out. Ensuing crystal structures were investigated after soaking at 20 minutes at sub-optimal concentration of PHB and optimal concentrations respectively. As seen in Figure 2A & 2B, PHB is captured by Aspartate 110 (Asp110) of one subunit by strong hydrogen bonding (2.32 Å) with –OH side chain of PHB while Arginine 12 (Arg12) along with Histidine 9 (His9) from other subunit aligns the ligand by several hydrogen bonds to carboxyl side chain. The orientation of PHB in dimer interface is maintained by the Arg12 and His9. The H-bonds with these two residues are not very strong (2.61 and 2.73 Å with Arg12; 2.94 Å with His9) as parallel molecular dynamics simulation (MDS) shows immediate release (and no rebinding) of PHB from these two residues in absence of binding Asp110 (from other subunit) in monomer structure (Supplementary movie 1). While MDS of monomer with Asp110 bound to PHB shows ligand in a continuous episode of binding and recapture in absence of supporting Arg12 and His9 from other subunit (Supplementary movie 2). The MDS of dimer reveals the presence of PHB in dimer interface bound to all three amino acids for some time before eventually departing from the cleft (Supplementary movie 3). There is a topspin rotational movement observed in the carboxyl side chain of PHB during these simulations (Supplementary movie 4). This top-spinning motion enables the ligand transitions from one subunit to the next within the dimer protein structure and ultimately departing from the protein. Additionally, several weak interactions during dimer simulation are also seen like opposing pi-pi interaction with Trp22 and Tyr34 and H-bonds with Ala35, Tyr34, Lys31 and Lys6 (from other subunit) (Supplementary movie 5).

**Fig. 2.**
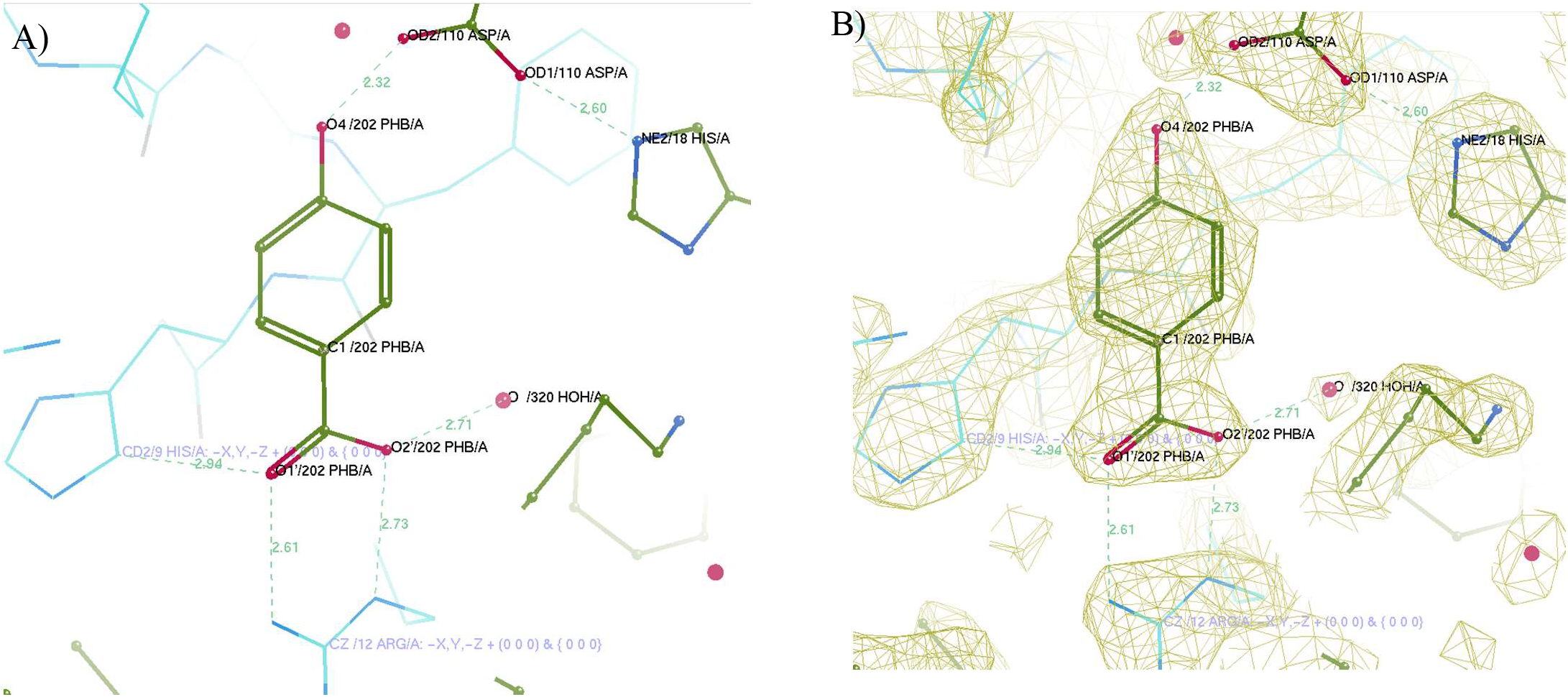
PHB interactions in HosA dimer interface. A) Model representation; B) Model along with 2Fo-Fc electron density map (pale green) contoured at 2σ. The main subunit is shown in dark green while the symmetry subunit in the dimer is shown in cyan in both A) and B). The bond distances in angstroms (Å) are shown as dashed lines (green) between PHB and respective interacting partners. The bond between the Asp110 from HosA and hydroxyl oxygen of PHB is the strongest and shortest one (2.32 Å). The other PHB interactions are with positively charged residues from symmetry subunit. These include two bonds with Arg12 (2.61 and 2.73 Å) and one bond with His9 (2.94 Å). These three interactions may play a role to align the Asp110 bound ligand in the HosA dimer interface. Apart from these, an association of PHB carboxyl oxygen (O2′) with a crystallographic water (320 HOH) is also shown (discussed later). The distance between Asp110 and neighbouring His18 is 2.6 Å in this structure. The figures are generated in COOT.

### The stability of the ligand in dimer interface is maintained by a water molecule (T3P11579) fluctuating in continuous tug-of-war among interacting amino acids from both subunits

Since, the MDS of PHB in absence of crystallographic water caused the ligand to be released from dimer cleft, we next focused on the experimental waters bound to PHB. There were totally 4 waters (three from same subunit, one from other subunit) bound to PHB (Figure 3A). Simulation carried out by keeping these waters in protein-PHB dimer exhibited utmost importance of the water (T3P11579) (Figure 3A) bound to PHB carboxyl side chain. The topspin rotation of PHB carboxyl side chain which was observed during previous simulations is abrogated in presence of this water (Supplementary movie 6). The other waters left very early during simulation (Supplementary movie 7). While T3P11579 along with PHB remained in position in dimer interface throughout entire 300 ns simulation (Supplementary movie 6). The structure showed T3P11579 interactions with PHB carboxyl side chain (while PHB bound to Asp110 with hydroxyl group on other side) and carbonyl oxygen of Phenyl alanine 8 (Phe8) from other subunit (Figure 3B & 3C), thus maintaining ligand in the dimer interface. Also, the close proximity of lysine 31 (Lys31) and Phenyl alanine 15 (Phe15) near PHB was observed in the structure (Figure 3B & 3C). MDS showed propeller like movement of Lys31 ε-amino side chain when it came close to T3P11579 (Supplementary movie 8). While the interacting hydrogen bond distances of PHB carboxyl group with Phe8 carbonyl group and T3P11579 were at synchrony, means they reduced or increased at same time points indicating characteristic attractive hydrogen bonding behaviours (Supplementary Figure 4A & 4B). On the contrary, there was no reduction in distance between Lys31 and T3P11579 with simulation time (Supplementary Figure 4C). This is characteristic of a repulsive interactions suggesting interactions between two species with similar charges. All these point towards the T3P11579 (also named as bridging water) as a positively charged hydronium ion (H_3_O^+^). In the structure, it is also interacting with Tyr34, Gly106 and Asp107 (all from same the subunit) demonstrating a propensity to bind to multiple partners owing to its excess valence. Further analysis by poseviewer script (poseviewer_interactions.py) as embedded in Schrodinger revealed several interactions with various other amino acids (from both subunits) with bridging water during simulation (Supplementary Table 2). Among these, most noteworthy were with Phenyl alanine 15 (from same subunit) and Tyrosine 34 (Tyr 34) from other subunit indicating pi-cation interactions. Throughout the simulation, the bridging water undergoes continuous fluctuations, driven by a constant tug-of-war among its interacting partners indicating dynamic nature of these interactions: most notably with Phe8, PHB, and Lys31 (Supplementary movie 8). This was also supported by the slightly higher B-factor (∼24) of this water in the crystal structure despite its proximity to several interacting partners.

**Fig. 3.**
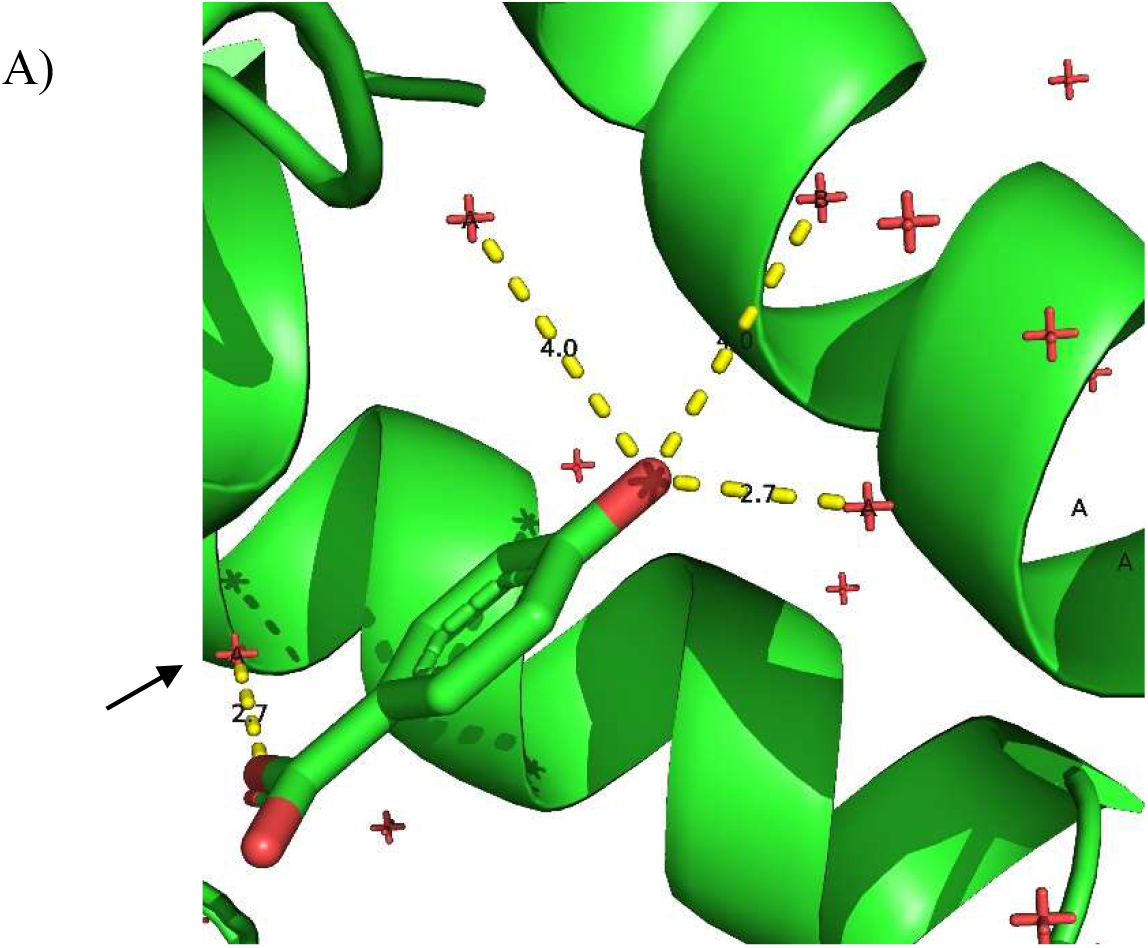
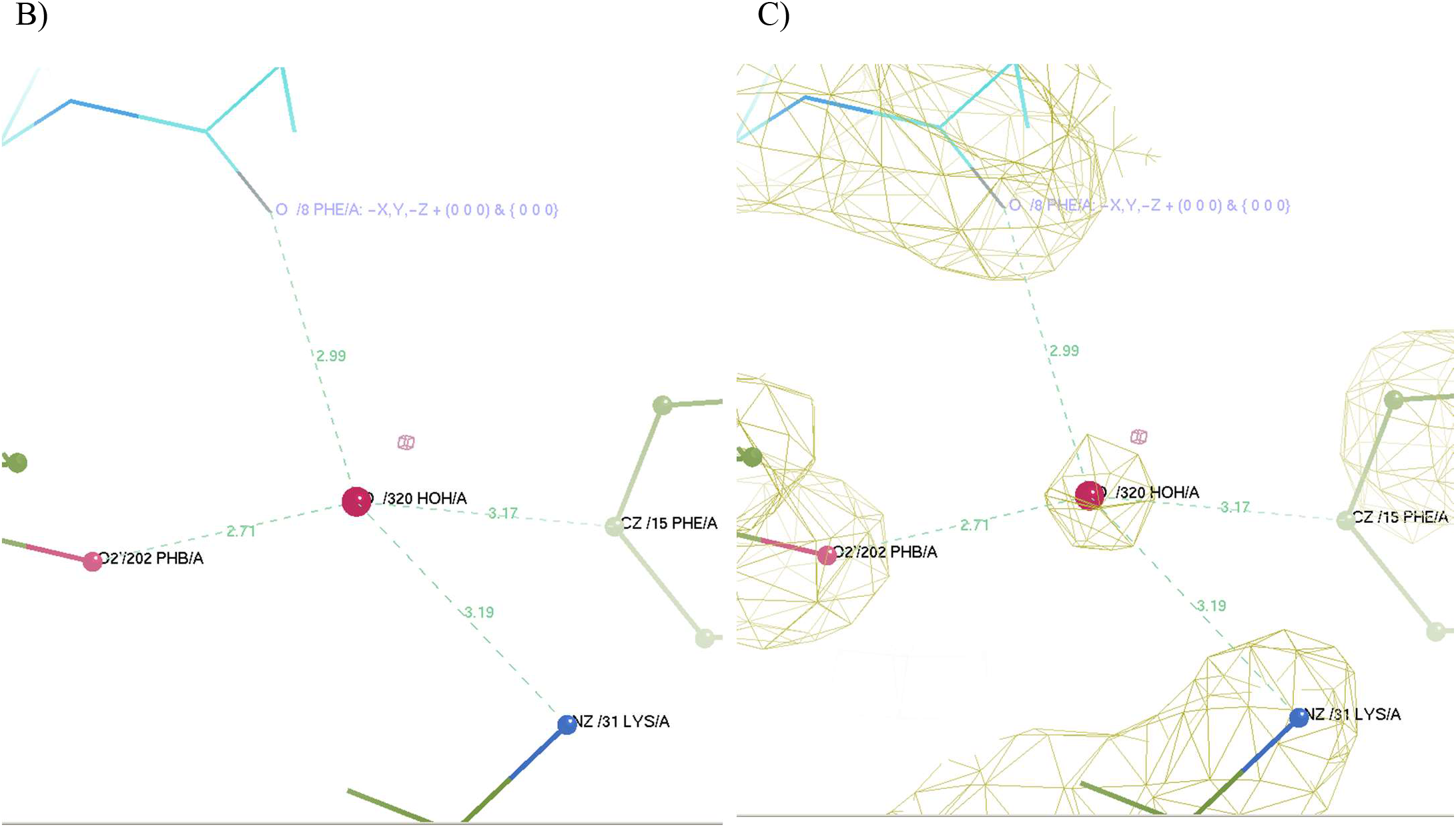
Bridging water for keeping PHB interactions in HosA dimer interface. A) Four crystallographic waters bound to PHB in HosA dimer generated by symmetry operation from 2.2 Å ligand bound structure. Bridging water (T3P11579) is shown by arrow. Three crystallographic waters including the T3P11579 are from the same subunit (denoted as A) and the remaining one is from the other subunit (denoted as B). The image is generated by PyMol (PyMOL-Molecular Graphics System, Open-Source version, Schrödinger, LLC). Except T3P11579, other waters left from the complex during initial steps of simulation. B) Model representation of 2.2 Å PHB bound dimer structure and C) along with 2Fo-Fc electron density map contoured at 2σ. The T3P11579 or bridging water is denoted as 320 HOH by the COOT generated images in B) and C). The bridging water network is shown by interacting Lys31, Phe15 and PHB from the same subunit chain (colored as green). The other interacting amino acid is Phe8 from symmetry subunit (colored in cyan). The corresponding interatomic distances (angstrom) are shown between the water and the interacting amino acids & PHB.

### Sequential involvement of two key histidines are responsible for subsequent ligand mediated conformational changes

From ligand soaked structure, the mechanism of PHB binding with the help of Asp110 in dimer interface and bridging water was elucidated. This resulted in PHB bound conformation I. Further, to explore later changes brought by PHB binding in dimer interface, co-crystallization experiments were performed. HosA protein was mixed with PHB at 1:2 ratio for crystallization. These crystals appeared after 7 days. The resulting 2.3 Å resolution structure revealed shrinking of the ionic bond between positively charged NE2 of Histidine 18 (His18) and negatively charged oxygen (OD1) from carboxyl side chain of Asp110 (Figure 4). Since the other carboxyl oxygen atom (OD2) from Asp110 remained bound to PHB, this juxtaposition of His18 is brought about by local conformational changes post ligand binding. The resulting inter-residual distance becomes 2.43 Å while it was 2.6 Å (Figure 4A & 4B) in the case of PHB bound conformation I. There was also clear enhancements of fused election densities from interacting atoms in the same bond path with 2Fo-Fc electron density map contoured at 2σ (Figure 4B). On the contrary, the 2Fo-Fc (2σ) electron densities of corresponding atoms were wide apart in ligand soaked structure (Figure 4A). Similar observation was noted for 2.16 Å resolution/diffraction limit structure solved via autoPROC-Staraniso-Phenix/Buster/Coot pipeline as well (Figure 4C). On the carboxyl side of PHB, Histidine 9 (His9) from other the subunit moves closer to the PHB O1′ atom which is away from the bridging water (Figure 4). There is also a clear rotation of His9 imidazole ring placing positively charged ND2 instead of CD2 (in ligand soaked structure) near negatively charged O1′ atom of PHB in co-crystallized structure (Figure 4B & 4C). The resulting attractive interaction distance shrinks from 2. 94 Å to 2.8 Å (2.75 Å in case of higher resolution structure solved through autoPROC-Staraniso-Phenix/Buster/Coot pipeline) (Figure 4B & 4C). This rotation and distance shrinkage are also observed in several structure solving trials of conformation II from multiple crystals (data not shown). In contrary, no such rotation was seen in His18 imidazole ring between the two structures. Parallel MDS of Apo-protein at 300 ns revealed the restricted mobility of the same being bound to Asp110 while solvent exposed His9 ring rotates freely in absence of ligand (Supplementary movie 9). Thus the enhanced proximity of respective histidines with Asp110 and PHB signifies the ligand binding mediated later conformational changes. This results in PHB bound conformation II of the dimeric protein which can bring about further downstream cascades.

**Fig. 4.**
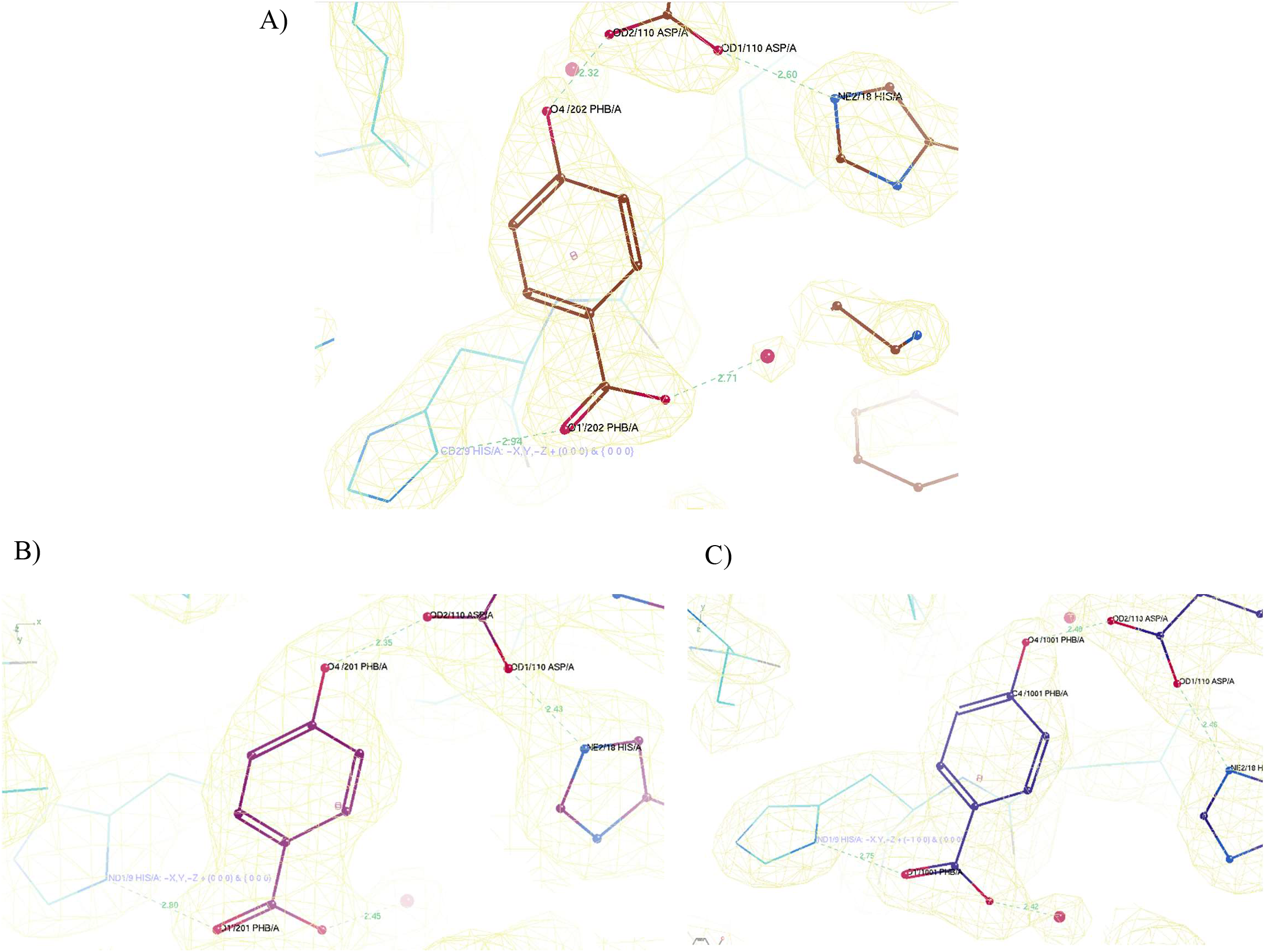
Role of histidine pairs post PHB binding in HosA dimer to result in ligand bound conformation II. The main subunit in the dimer is colored as brown (A), red (B) and blue (C) while the symmetry subunit as cyan. A) Initial conformation I (2.2 Å resolution structure) after PHB binding has His18 wide apart from ligand bound Asp110 with an inter-residual distance of 2.6 Å. The electron densities of the respective amino acids are also remain unmerged. The carboxyl side chain of PHB is far from the neighbouring His 9 from the other subunit (generated by symmetry operation) in the protein dimer. The distance between PHB carboxyl -O and nearby -C_δ_ (or –CD) of imidazole ring of His9 is 2.94 Å. B) His18 juxtaposes to Asp110 in PHB bound conformation II (2.3 Å resolution structure), with an inter-residual distance of 2.43 Å and clear fusion of respective residual electron densities. The His9 also rotates to result in shrinkage of distance with PHB to 2.8 Å via placing positively charged -N_δ_ (or –ND) of imidazole ring near negatively charged oxygen of PHB carboxylic side chain. C) The respective His18 and His9 show same behaviour at 2.16 Å resolution structure as well (autoPROC-Staraniso-Phenix/Buster/Coot pipeline). The distance shrinks to 2.46 Å in the former while it reduces to 2.75 Å in the latter. In all the COOT diagrams, the 2Fo-Fc densities (pale yellow) are contoured at 2σ.

### The ligand bound T3P11579 mediated subtle changes in dimer interface happens via flipping of Phenyl alanine 15 (Phe15) from both subunits

One of the most important interactions in HosA apoprotein dimer interfaces are formed by Phenyl alanine 15 (Phe15) from both subunits. Simulation of the same reveals continuous vigorous flipping of Phe15 aromatic rings from both subunits, interrupted periodically by transient interactions with Lys31 (from both subunits) and sometimes Phe8 (from one subunit only) (Supplementary movies 10 and 11). After binding to PHB in conformation I, this flipping is changed to partial rotation, indicating influence from bridging water T3P11579 (attached to PHB), to affect more the nearer Phe15 than the distant one (Supplementary movie 12). Major changes are observed between ligand bound conformations I and conformation II. A distinct rotation was observed in Phe15 from both subunits when these two structures are superposed (Figure 5A & 5B). These rotations are observed in conformation II from both 2.3 Å (Figure 5A) and 2.16 Å (Figure 5B) resolution structures. The critical positioning of Phe15 pairs marks the location of the hinge region, situated between the dimerization tip and the DNA binding lobes (Supplementary Fig. 3C) of crown shaped HosA dimer. Any changes in this region can disturb proper functioning of DNA binding lobes located beneath as shown for other MarR structures. Interestingly, the histidine pairs (His9 and His18) as discussed previously are also located in this region only. Therefore, it can be speculated that the striking rotations of dual Phe15 residues, histidine pairs’ movements and the bridging water associated amino acids drive the conformational changes in the hinge region, necessary for the DNA dissociation in PHB bound conformation II.

**Fig. 5.**
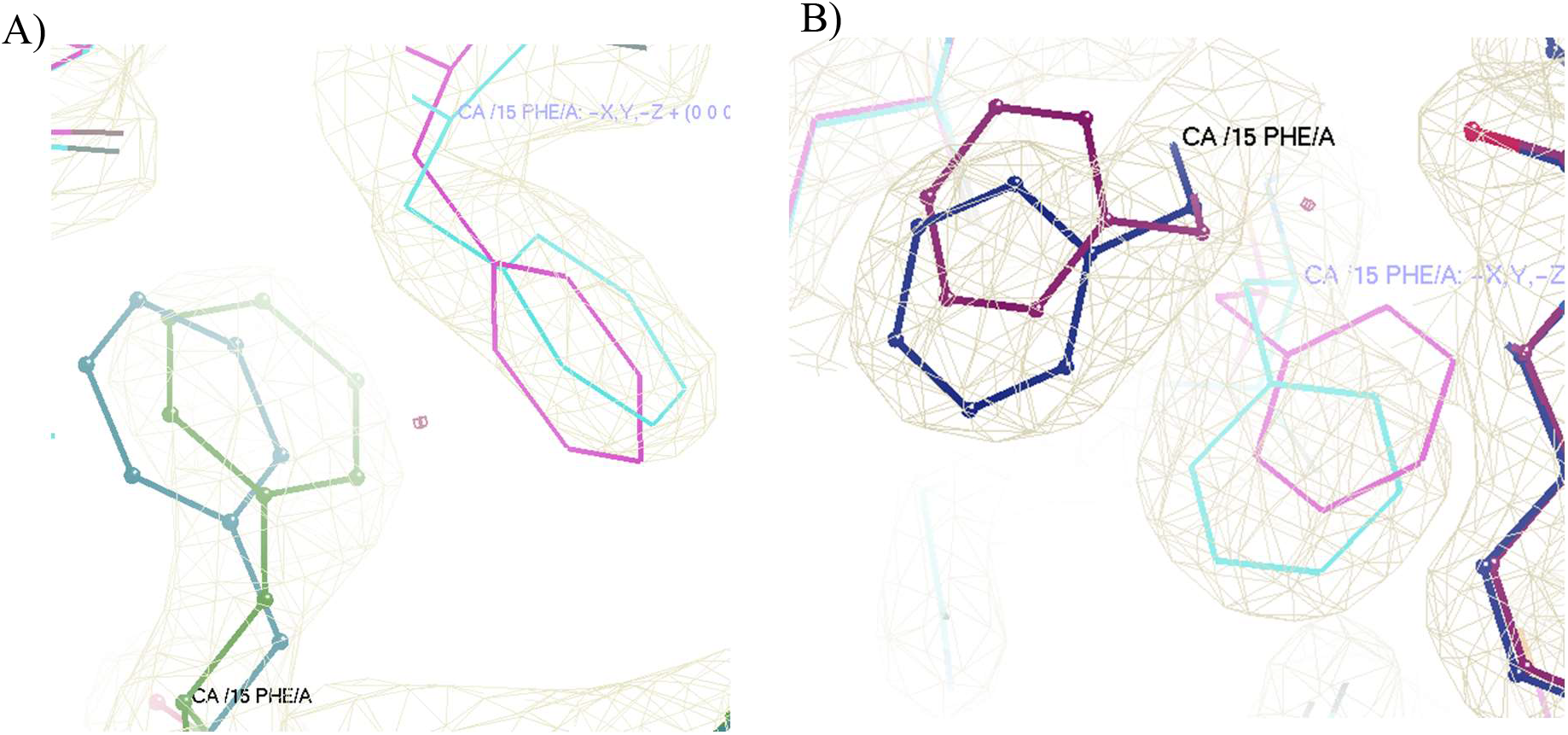
Synchronous rotations are observed in the Phe15 from both subunits in PHB bound conformation II. A) As dimers, PHB bound conformation II (2.3 Å resolution structure) is aligned on conformation I (2.2 Å resolution structure). The distinct rotations are seen in Phe15 residues from both subunits. The other subunit is generated by symmetry operation in the protein dimer. The conformation I residues (from the main subunit) are colored as blue and conformation II residues (from the main subunit) are colored as green. The symmetry molecules of these structures are shown in pink and cyan respectively. B) Similar clear rotation is noted for Phe15 pairs in 2.16 Å resolution dimer structure as well (autoPROC-Staraniso-Phenix/Buster/Coot pipeline) aligned on conformation I. The conformation I residues (from the main subunit) are colored as red and conformation II residues (from the main subunit) are colored as blue. The symmetry molecules of these structures are shown in pink and cyan respectively. In both COOT diagrams, the 2Fo-Fc densities (pale green) are contoured at 2σ.

## Discussions

Once upon a time there were two bacteria exchanging genetic material in sewage water. One of them was pathogenic *Escherichia coli*, another one was *Shigella*. That was the crucial time point enteropathogenic *E coli* got HosA gene from *Shigella*. The end result is further evolutionary shifting of pathogenic *E. coli* from non-pathogenic ones due to presence of one more class of MarR type proteins in the former ones. Particularly, this also enabled these *E. coli* to metabolize benzoic acid compounds present in sewage discharges. Interestingly, shiga toxin gene was also received by horizontal transfer (bacteriophage mediated) which led to development of Shiga toxin-producing *Escherichia coli* (32). Thus horizontal transfer has shaped evolution of not only antibiotic resistance, but development of toxin producing and chemical resistant strains. The entire operonic sequence of HosA got transferred enabling the recipient *E. coli* to produce all enzymes required for non-oxidative metabolism of benzoic acid compounds such as PHB by scavenging from sewage. The significance of this metabolism is clear from the presence of HosA gene in almost all pathogenic *E. coli strains* according to Uniprot (33).

The stable binding of PHB to HosA is orchestrated by a single water bound to its carboxylic group. The significance of the water is revealed by extensive molecular dynamics simulation showing essentiality of the water to bridge individual subunits in the dimer as well as to bring about the shuffling of residue side chains to result in disturbances in the dimer interface. This may cause the DNA to be released from the dimer. So instead of large scale energy exhausting conformational changes involving dimer to monomer transitions to be released from DNA, HosA applies subtle changes in the dimer interface to bring about the same outcome in an energy efficient manner. This approach was previously speculated for MarR transcription factors as large conformational changes were observed upon DNA displacing ligand binding in the hinge region located in the dimer interface (3, 34) the details of which was not explored further. Our study shows the detailed mechanism beneath these alterations in the hinge region by specifically displaying specific amino acid movements through extensive molecular dynamics simulations in real time. Particularly, Phe 15 rotations from both subunits and His9-His18 coupled movements are elucidated. Further, the role of highly fluctuating water and associated amino acids as the bridging entity between PHB and protein as well as between protein subunits is elucidated.

The bridging water (T3P11579) is in a highly flexible position connected by all three sides in a tug-of-war machinery by ε -amino group of lysine (Lys31), Phenyl alanine (Phe8) carbonyl (from other subunit) and PHB carboxyl group. Apart from this, we also identified several more interactions with other amino acids (from poseviewer_interactions script analysis), specially with phenyl group of Phe15 (Supplementary Table 2). This indicates the water to be a hydronium ion (H3O^+^) having cation-pi interaction with phenyl ring of Phe15. This also explains the positional fluctuations in the water being an ionic entity with extra valence from additional oxygen, ready to interact with several interacting partners. Further analysis from distance vs. simulation time plots of the water and interacting residues reveal that the interactions between water and Phe8 or PHB are attractive ones as opposite to the repulsive interaction with Lys31. There was significant reduction in distances between T3P11579 and corresponding interacting atoms of either Phe8 or PHB, particularly within 100-140 ns range before increasing again (Supplementary Figure 4A & 4B). This is indicative of attractive ionic interaction, in which oppositely charged ions come closer to each other until there is steric hindrance from their respective Vander Waals’s radii and neighbouring atoms (35). All interacting partners from either Phe8 or Phe15 or PHB are negatively charged (Fig. 3B) indicating O of T3P11579 as positively charged. In case of repulsive interactions, the interatomic distances cannot shrink significantly, rather remain constant owing to interacting mobile atom pairs’ and neighbouring atoms’ Vander Waal’s contact distance limits. This was seen in the bond distances between Lys31 positively charged imino H and O of T3P11579 which remained unchanged during the simulation (Supplementary Figure 4C). Therefore, the O of T3P11579 may be positively charged as in a hydronium ion (H3O^+^). Further atomic resolution diffraction is needed to support this property of the bridging water.

## Acknowledgements

AG thanks Rukmini R and Mallesham K for technical support during data collection in X-ray facility, CSIR-CCMB, Hyderabad, India. Also, Dr. Kavyashree Manjunath, InStem, Bangalore, India is duly acknowledged for her suggestions during data processing and synchrotron data collection. The contributions by the staffs of PX-BL21, Indus-2 is suitably acknowledged during synchrotron data collection. AG also acknowledges the riveting discussions on ccp4bb bulletin titled “About Staraniso” on Mon, 12 Feb 2024 for significant insights.

## Author contributions

AG: Conceptualization, Resources, Experiments design & performing, Structure solving, Analysis, and Writing. SU: Molecular dynamic simulation. JAB: Structure solving by autoPROC-Staraniso-Phenix/Buster/Coot pipeline.

