## Supplementary Fig. 1-4 for "Horizontally acquired HosA transcription factor bound with 4-hydroxy-benzoic acid exhibits unique tug-of-water dynamics"

**Supplementary Fig 1. Sequential alignment of HosA**
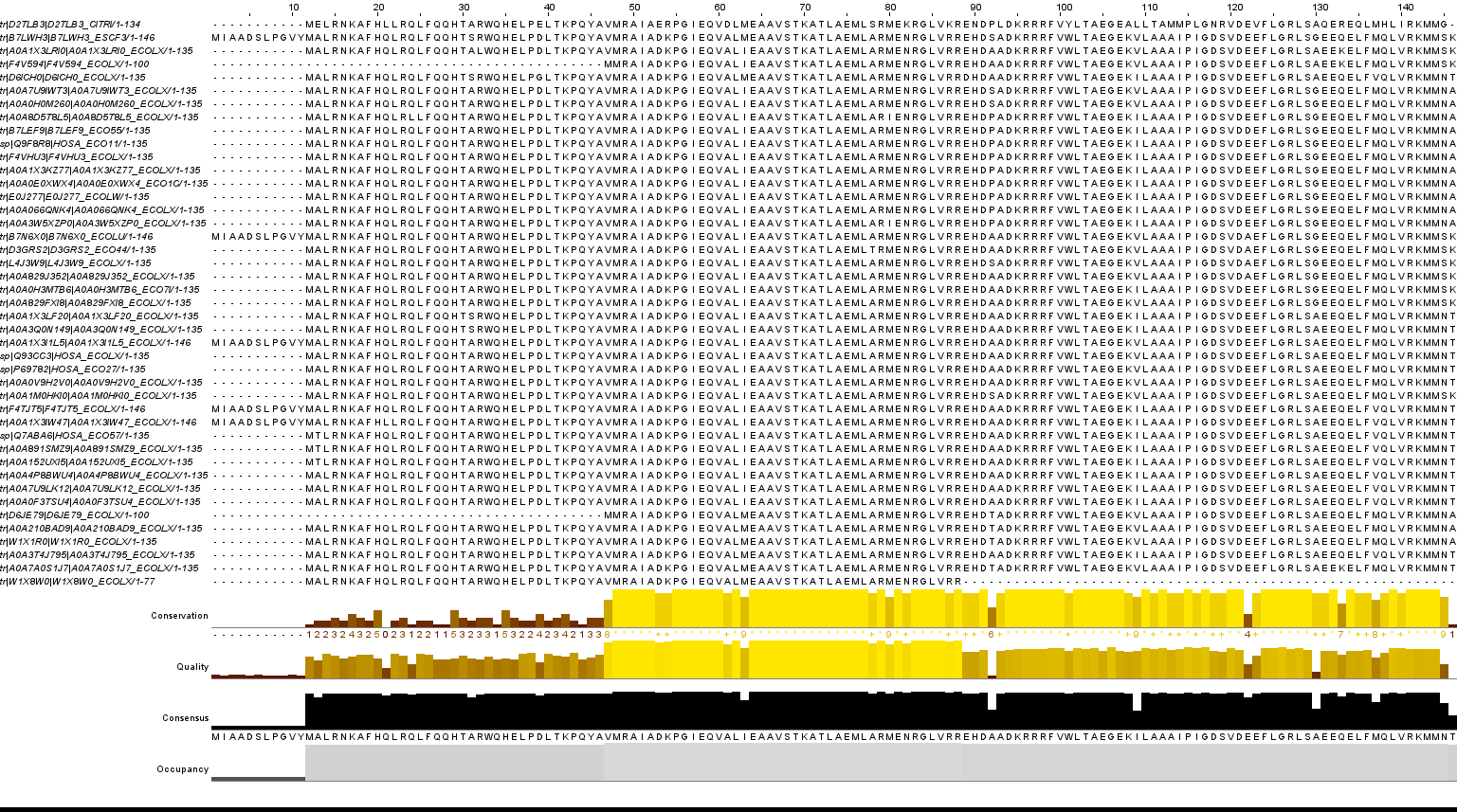
**across pathogenic *E. coli* homologs.** As shown in bottom bars, high sequence conservation was noted among these sequences (>70% identity). The sequence alignment is from Clustal omega (1). The image is generated by Jalview (2).

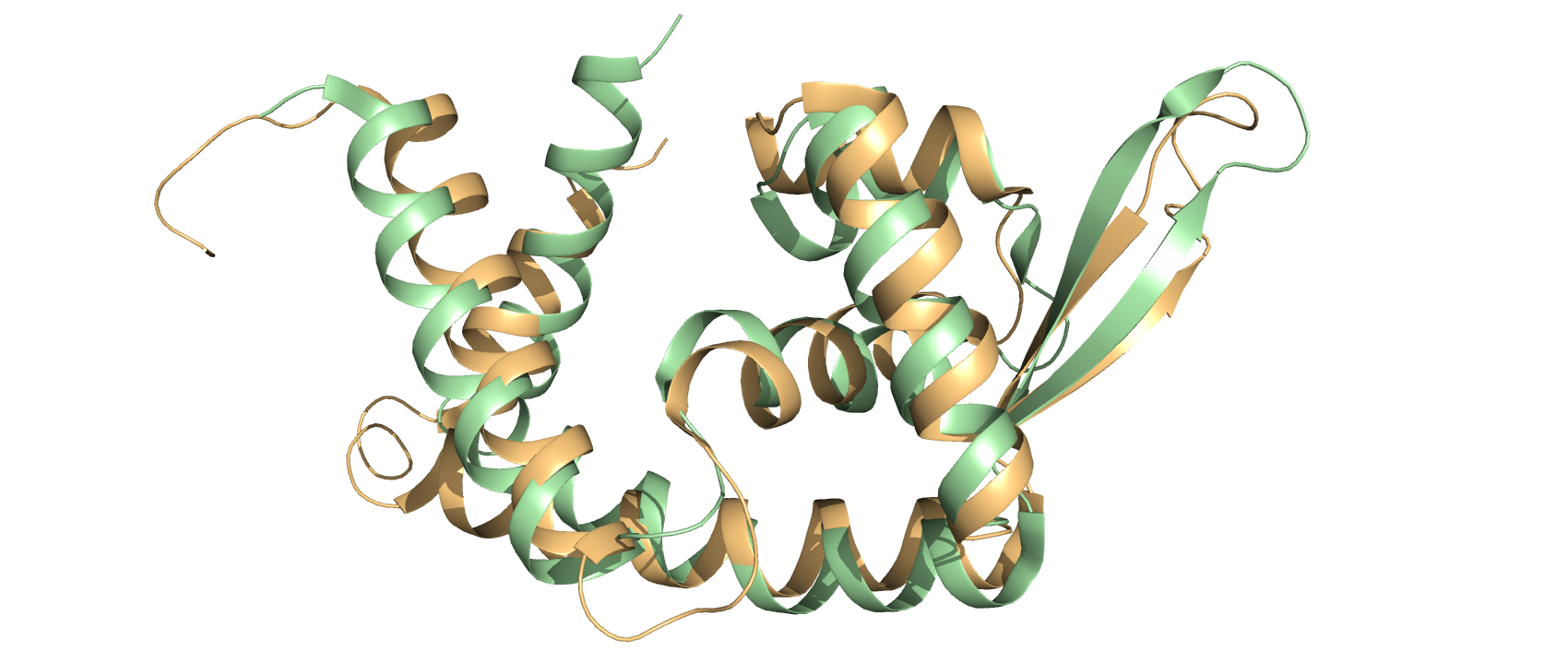

**Supplementary Fig 2. Structural alignment of EPEC HosA on *E. coli* MarR.** Structural alignment of HosA apoprotein (green) (8PQ4) on *E. coli* MarR (orange) (1JGS) displays high degree of similarity with similar structural motifs. The distinct local kink on winged β-sheet of HosA is marked by the arrow. The image is generated by PyMol (PyMOL-Molecular Graphics System, Open-Source version, Schrödinger, LLC).

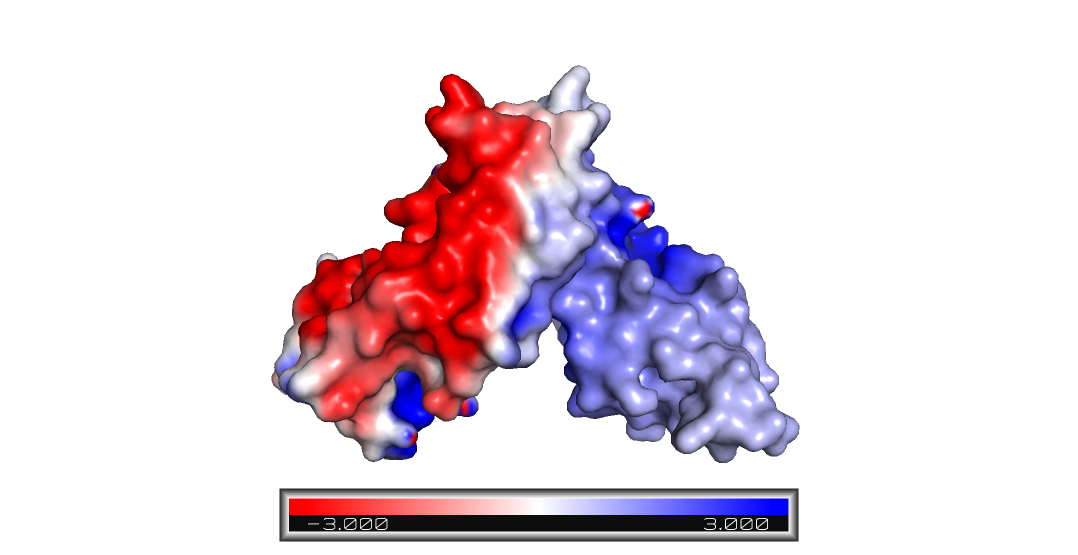

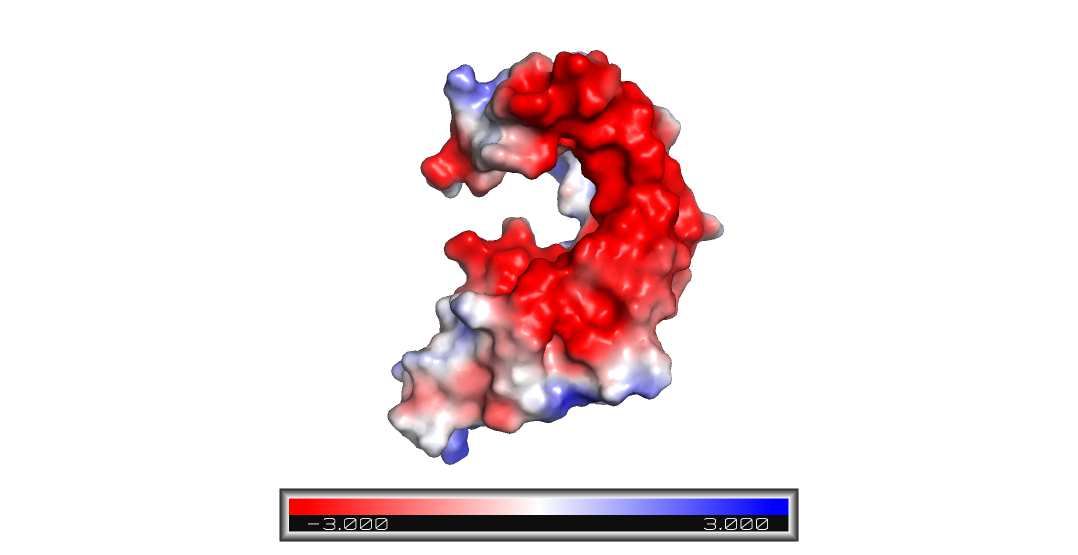

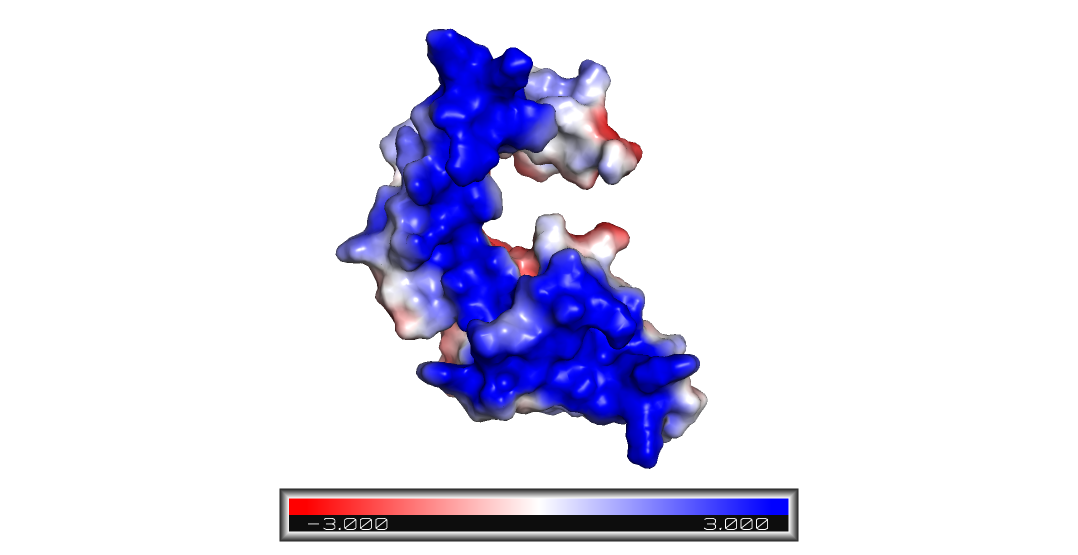

**C)**

**B)**

**A)**

Hinge

**Supplementary Fig 3. PDBePISA and APBS analysis shows HosA as a dimer formed by associations via oppositely charged surfaces of individual monomers.** Monomer with **A)** negatively charged surface (red) and **B)** positively charged surface (blue) at association point. **C)** The resulting crown shaped dimer has abundant positive charges at bottom surface, ready to interact with negatively charged DNA. The electrostatic charges are represented by red-white-blue color bar (-3.000 to +3.000). The 2-fold rotation axis for the dimer is shown by arrow and dashed line. The image is generated by PyMol.

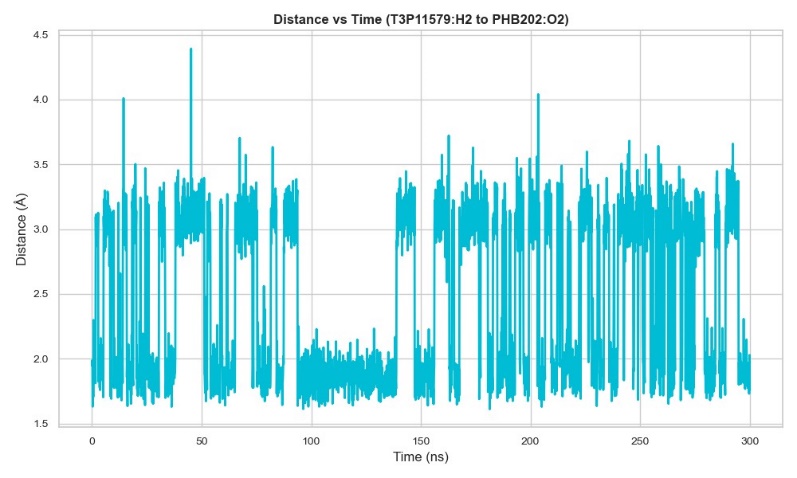

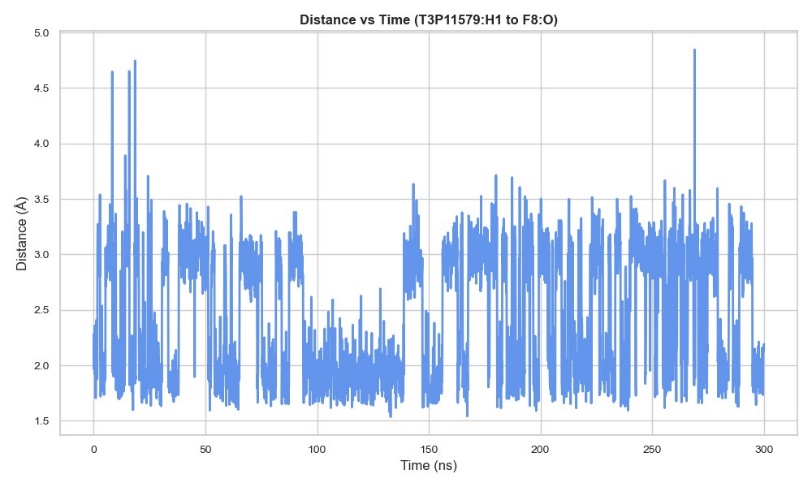

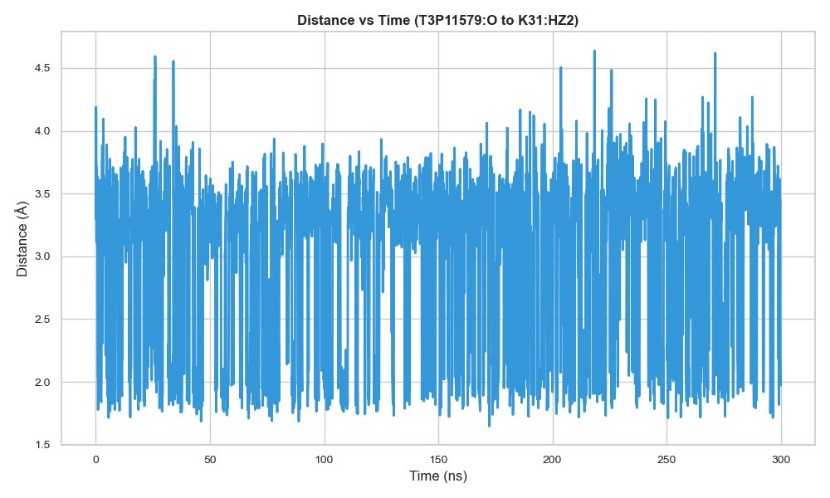

C)

B)

A)

**Supplementary Fig 4. X-Y plots of time dependent dynamic distance variations between bridging water and interacting partners indicate both attractive and repulsive ionic forces at play.** Dynamic distances (Å) between bridging water (T3P11579) and interacting HosA dimer residues & PHB are generated with respect to time (ns) from 300 ns simulation. The closest interatomic distances are plotted. A) Distance variations between T3P11579 hydrogen and PHB carboxyl oxygen vs. time (ns) are shown. B) Similar time dependent distance changes between T3P11579 oxygen and Phe8 (F8) carbonyl oxygen are observed. Most notably, almost identical reduction of interacting distances are observed within 100 to 140 ns in both A) and B). This type of profile indicates attractive ionic interactions where connecting pairs tend to come closer for significant period of time until blocked by steric hindrances from Vander Waals’s forces. C) Absolutely no significant time dependent reduction of interacting distances between T3P11579 oxygen and Lys 31 (K31) imino hydrogen is observed. This indicates a repulsive interaction formed by similarly charged atoms, maintained by almost invariable interacting distances as controlled by Vander Waals’s radii. Transient changes (up and down) can be seen in the dynamic distance trace as expected from two interacting but continuously mobile atoms.

**Supplementary Methods and Results**

**Molecular dynamics methodology**

**Simulation system preparation and trajectory production**

The HosA-apo and HosA-p-hydroxy benzoic acid complex were imported into the desmond simulation algorithm for system building (3). Both the HosA apo and ligand bound structures (both apo and dimer structures) were solvated in a TIP3P (transferable intermolecular potential 3P) water model (4). The orthorhombic box was selected, and the boundary was set to 10 Å in both cases. A salt concentration of 0.1M (molar) was added in both cases and the neutralization of the system achieved by adding Na ions. The full system contained 22286 atoms in case of HosA-apo and 24869 atoms in case HosA-p-hydroxy benzoic acid complex. The NPT ensemble class with Temperature 300 K and pressure 1 atm was used to simulate each system for 100 ns and 300 ns separately using MD (Molecular Dynamics). The OPLS2005 force field and the default "relax model before simulation algorithm" protocol were selected before trajectory production (5). The simulation results for both systems were analysed for (root mean square deviation-RSMD), (root mean square fluctuation-RMSF), (radius of gyration-Rg), (solvent accessible surface area-SASA), (polar surface area-PSA), and (hydrogen bonds-HBonds for complex). Lastly, the simulation quality analysis was conducted for each system. The AMD**^®^**^TM^ Ryzen 7 CPU (*central processing unit*) with 64-GB of RAM on Linux Fedora 37 Scientific OS (operating system) was used for the simulations and the analysis.

**Supplementary molecular dynamics results**

**Conformational Dynamics and Stability of monomer HosA- p-Hydroxy Benzoic Acid in ligand soaked structure (Conformation I)**

We conducted a detailed analysis of the dynamics and stability of monomer HosA in its apo form and in complex with p-hydroxy benzoic acid (8AGA) using 100ns trajectories. The rmsd graph of the HosA-apo structure simulation revealed a highly stable structure with low fluctuations, as indicated by the maximum rmsd of 4.65 Å at 18.10ns and an average rmsd of 3.86 Å. The histogram of the apo structure further confirmed this trend, with a consistent distribution of conformations throughout the simulation (Figure 1B). In contrast, the HosA-p-hydroxy benzoic acid complex demonstrated a more dynamic and less stable structure, with a higher rmsd of 7.13 Å at 98.90 ns and an average of 5.91 Å at the 100ns trajectory (Fig. 1C-D). To gain a deeper understanding of these findings, we extracted frames from both trajectories and superposed them on the starting frame of 0ns as a reference. The superposition of the HosA-apo structure at 18.10 ns revealed fluctuations in the N-terminus region residues (ASN1 to HIS5), centre vicinity region (VAL45 to ALA50), and the C-terminal region (ALA115 to ASN130) (Figure 1E-G). These fluctuations suggest that these regions are more flexible and might play an important role in the dynamics of the protein.

**Ligand Displacement in HosA Complex**

Interestingly, the superposition of the HosA-p-hydroxy benzoic acid complex at 98.90ns revealed a significant displacement of the ligand from its parent centre vicinity towards the N-terminus. The p-hydroxybenzoic acid formed a network of four hydrogen bonds with HosA, two with ARG9 at distances of 2.24 Å and 2.09 Å, one with THR16 at 1.88 Å and another with GLN13 at 2.29 Å (as shown in Figure 2A). This shift was first observed at 1.20ns and was further confirmed in the (supplementary Movie1), where the ligand moved towards the N-terminus and remained in that region for the entire 100ns simulation trajectory in Figure 2B. Furthermore, to better understand the displaced position and the time spent by the p-hydroxybenzoic acid was captured in a times-based gradient representations of full trajectory at each 10^th^ ns frame in contact with the HosA structure at the N-terminus region (as shown in Figure 1D).

**HosA-p-Hydroxy Benzoic Acid Complex is More Compact and Tightly Packed**

In figure 3A-B, the radius of gyration was determined for HosA in its apo form and its complex with p-hydroxybenzoic acid. The average value of the radius of gyration for HosA-apo was found to be 18.04 Å, whereas the average value for HosA-complex was 15.82 Å. This suggests that the complex is more compact, tightly packed, and exhibits a well-defined interface between HosA and the ligand (supplementary Movie2). The highest observed radius of gyration value for HosA-apo was 18.89 Å, while for the complex it was 20.94 Å. These findings suggest that the binding of p-hydroxybenzoic acid induces a conformational change in HosA that results in a more compact structure with a tighter interface. In Figure 4A-D, the root mean square fluctuation (RMSF) analysis of HosA protein in both its apo and complex conformations was conducted to investigate the protein's structural flexibility and rigidity. The average RMSF value recorded for HosA-apo was 1.93 Å, indicating that the protein is relatively flexible in its unbound state. Furthermore, the RMSF analysis revealed that the C and N-terminus regions of HosA-apo were more flexible compared to the central vicinity. In contrast, the RMSF analysis of HosA-complex revealed that the N-terminus and C-terminus region of the protein was observed to be more flexible with an average RMSF value of 1.66 Å, while the central remained relatively rigid. This suggests that the binding of the protein to its ligand induces a conformational change in HosA, resulting in a more stable protein-ligand complex at N-terminus rather than in the vicinity or central region. The analysis of the HosA protein in its unbound (apo) and bound (complex) states revealed significant differences in its solvent accessible surface area (SASA) values. Specifically, the highest SASA value observed for HosA-apo was 9569 Å2, with an average value of 8737 Å2. In contrast, for HosA-complex, the highest SASA value was 9440 Å2, with an average value of 8250 Å2 (as presented in Figure 5A-B). These observations suggest that the binding of p-hydroxy benzoic acid induces significant structural changes in HosA, resulting in a more compact conformation with decreased accessibility to solvent molecules. The decrease in SASA values upon ligand binding is consistent with the root mean square deviation (rmsd) and radius of gyration (rg) observations, further supporting the notion that the binding event causes significant conformational changes in HosA.

**Hydrogen bond analysis of HosA-p-Hydroxy Benzoic Acid Complex**

A detailed analysis of the hydrogen bonding interaction between HosA and p-hydroxy benzoic acid over a 100 ns trajectory conducted, which revealed crucial insights into the key interactions between HosA and p-hydroxy benzoic acid. In particular, we observed a higher frequency of hydrogen bonds formed between p-hydroxy benzoic acid and HosA residues ARG12, GLN16, and THR19, as depicted in figure 6A-C. Furthermore, the trajectory analysis also revealed the presence of protein internal hydrogen bonds, as illustrated in figure 6D. These results further provide a more understanding of the hydrogen bonding interaction between HosA and p-hydroxy benzoic acid and highlights the specific amino acid residues that are involved in stabilizing this interaction at 100ns simulation.

**Figure 1B** The histogram illustrates the frequency distribution of RMSD values for the HosA apo structure. The green-colored dashed line corresponds to the maximum RMSD value of 4.65 Å, while the pink-colored dashed line corresponds to the average RMSD value of 3.86 Å. A time-based color gradient from blue color corresponds to 0ns to red 100ns can also be observed in terms of HosA-apo structure fluctuations.

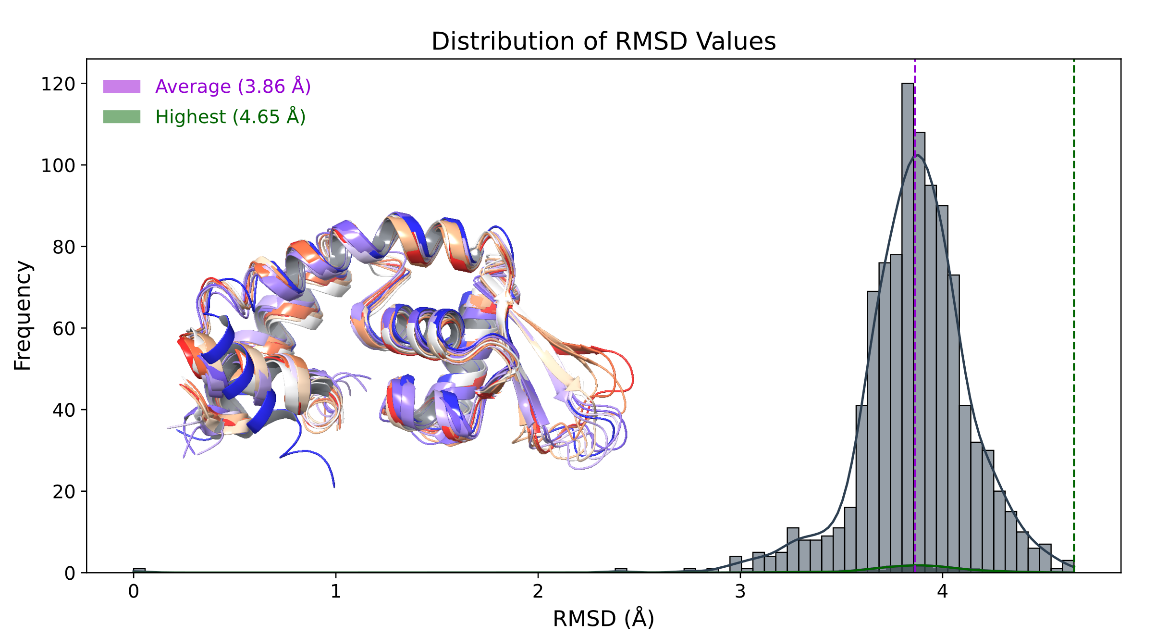

**Figure 1A** displays the Root Mean Square Deviation (RMSD) of the HosA-apo structure during a 100ns simulation. The black arrow points to the highest RMSD value of 4.65 Å, which occurs at 18.10ns, while the average RMSD value of 3.86 Å is represented by the dashed gray line.

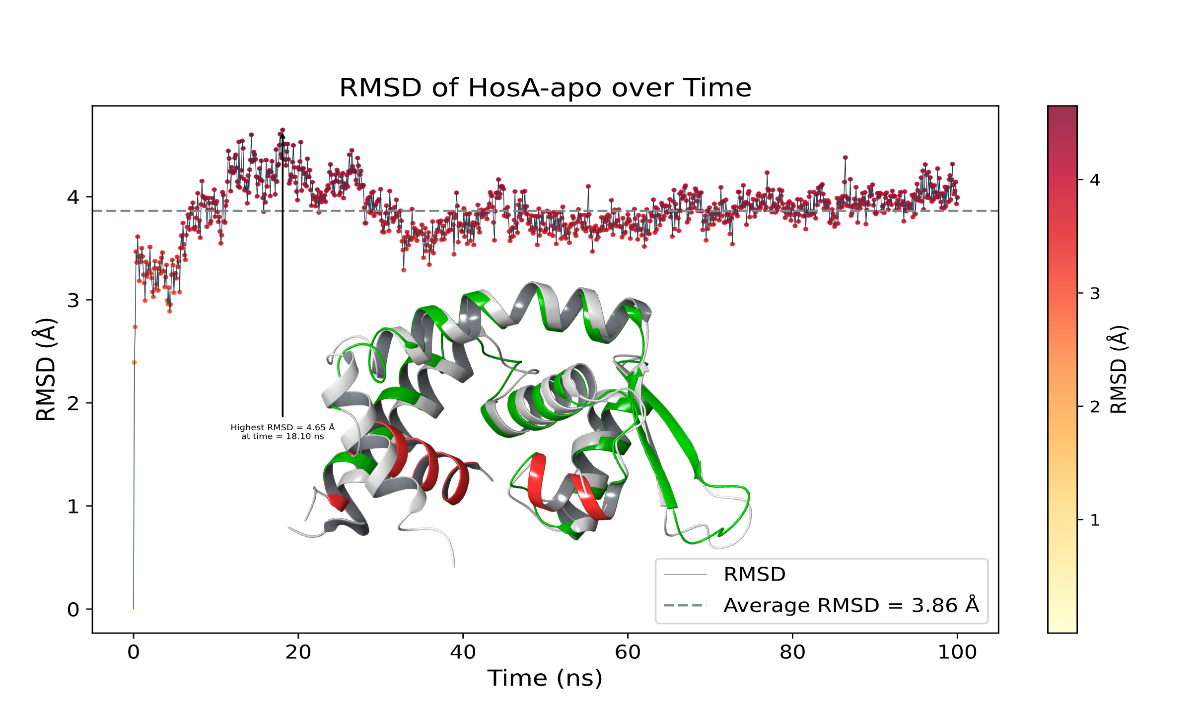

**Figure 1C** Highest structural fluctuation observed in HosA-p-hydroxybenzoic acid complex at 98.90ns with rmsd of 7.13 Å (denoted by black arrow). Average rmsd (5.91 Å) indicated by light gray dashed line. Green superposed structure of complex at 98.90ns and pink mesh of ligand at 0ns for comparison

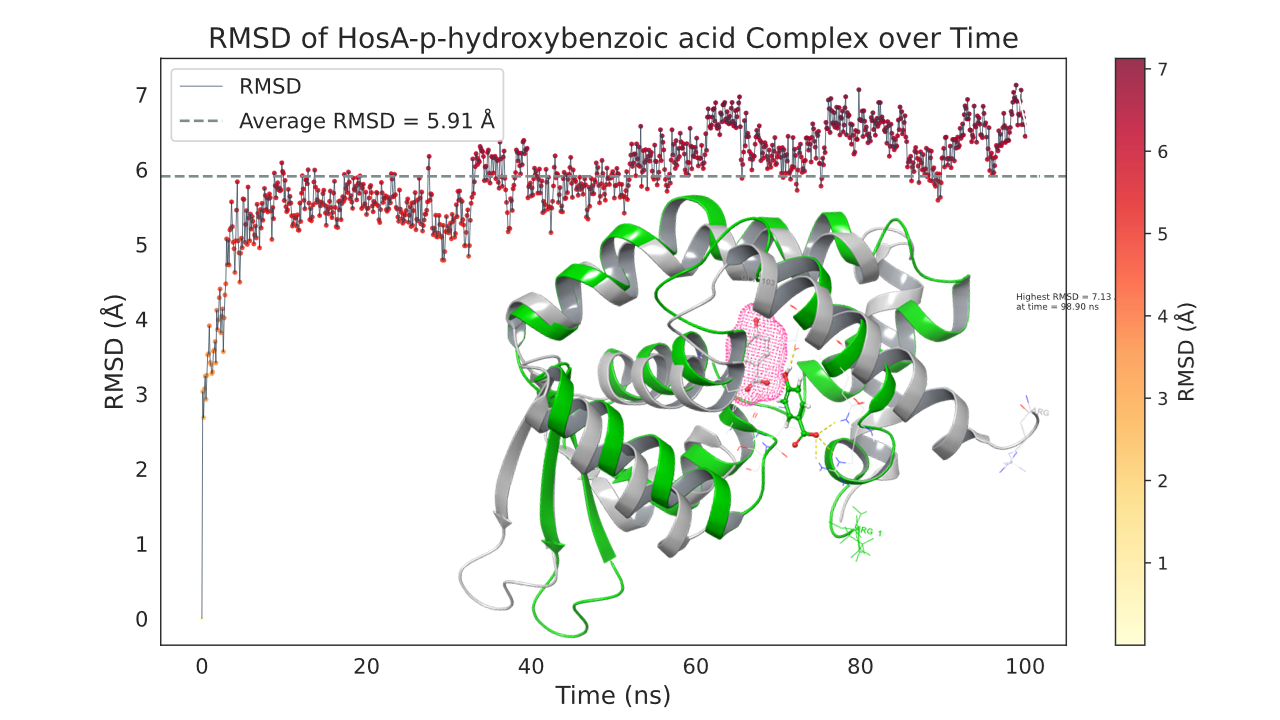

**Figure 1D** The histogram distribution of root-mean-square deviation (RMSD) for the HosA-p-hydroxybenzoic acid complex shows that the highest value of 7.13 Å represented by a green dash line. The average value of RMSD, represented by a pink dash line, is 5.91 Å. A time-based color gradient ranging from blue (0ns) to red (100ns) is noticeable for p-hydroxybenzoic acid.

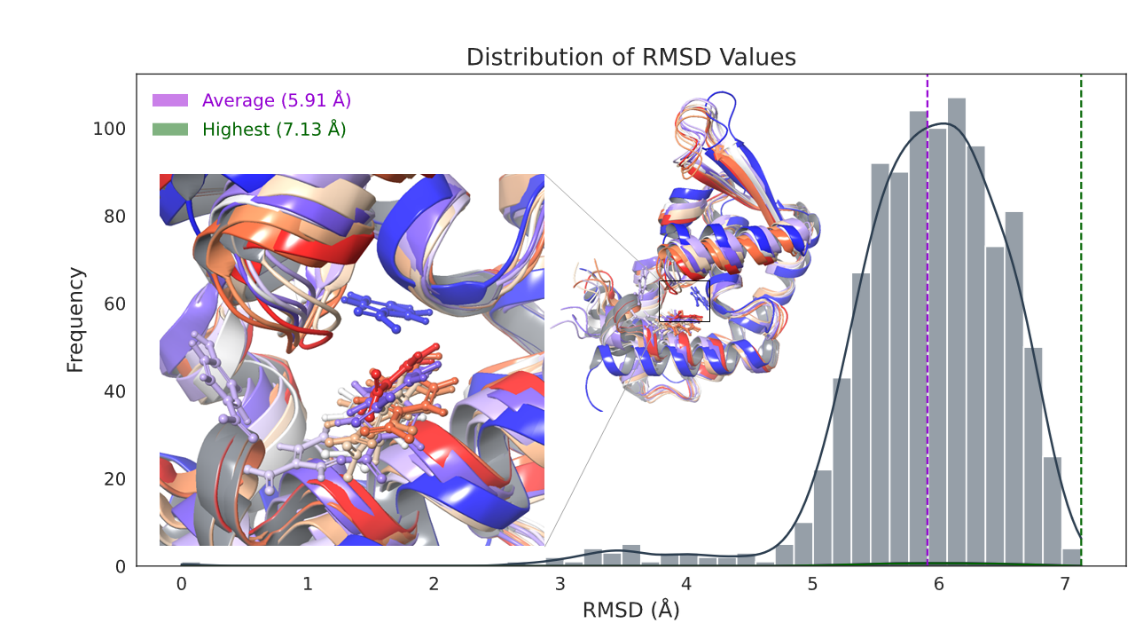

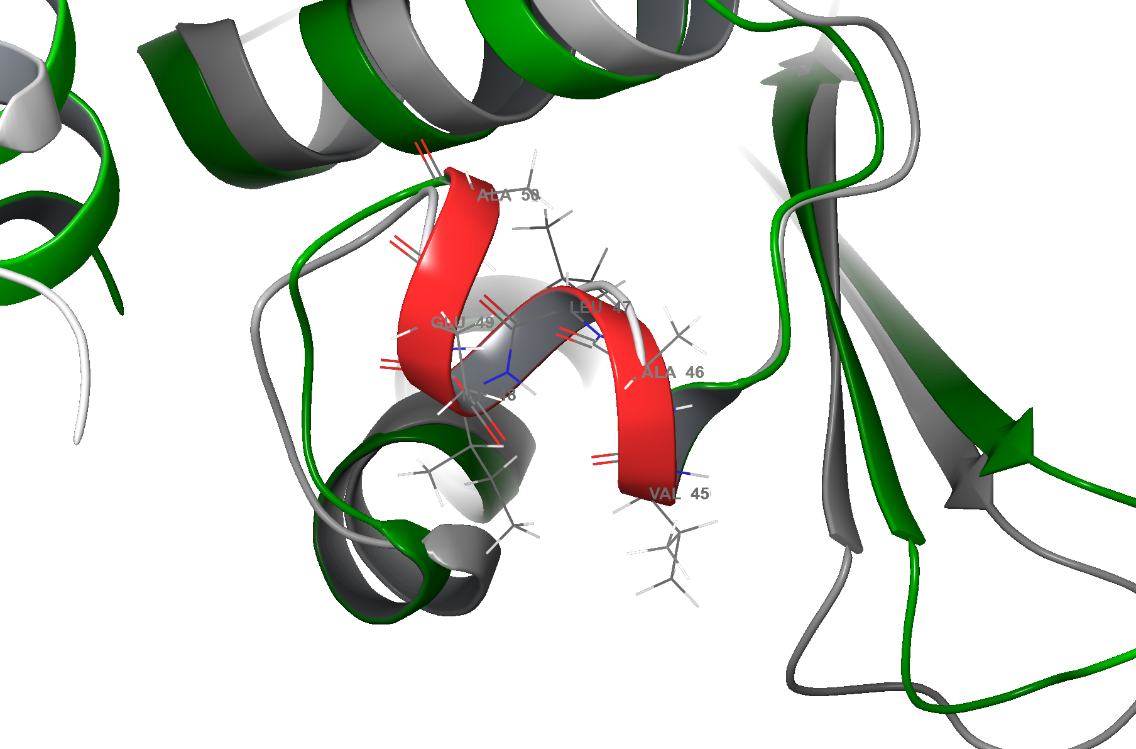

**Figure 1F** At 18.10ns, the center loop region of HosA-apo structure is depicted in green and structural fluctuations in red, superimposed on the 0ns reference frame in grey color.

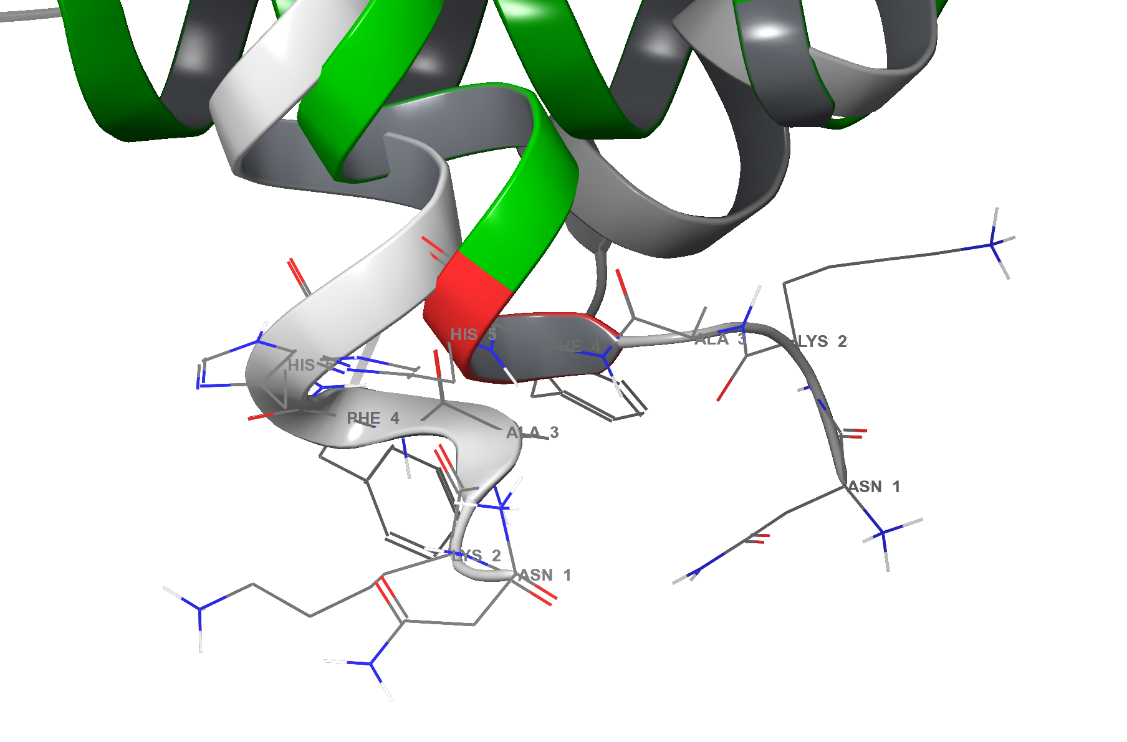

**Figure 1E** The HosA-apo structure's N-terminal region is shown in green with red structural fluctuations at 18.10ns, overlaying the grey 0ns reference frame.

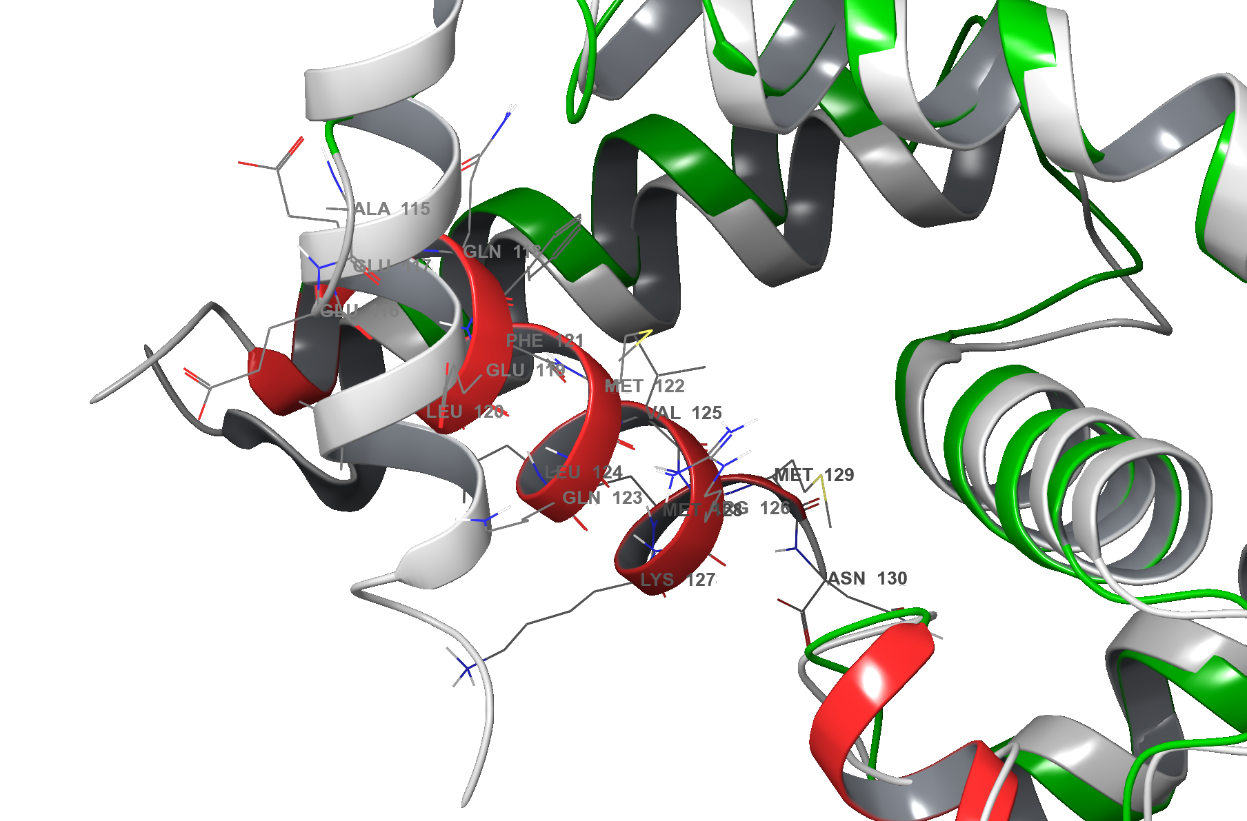

**Figure 1G** The HosA-apo structure's C-terminus (green) and structural fluctuations (red) at 18.10ns are shown overlaid on a grey reference frame at 0ns.

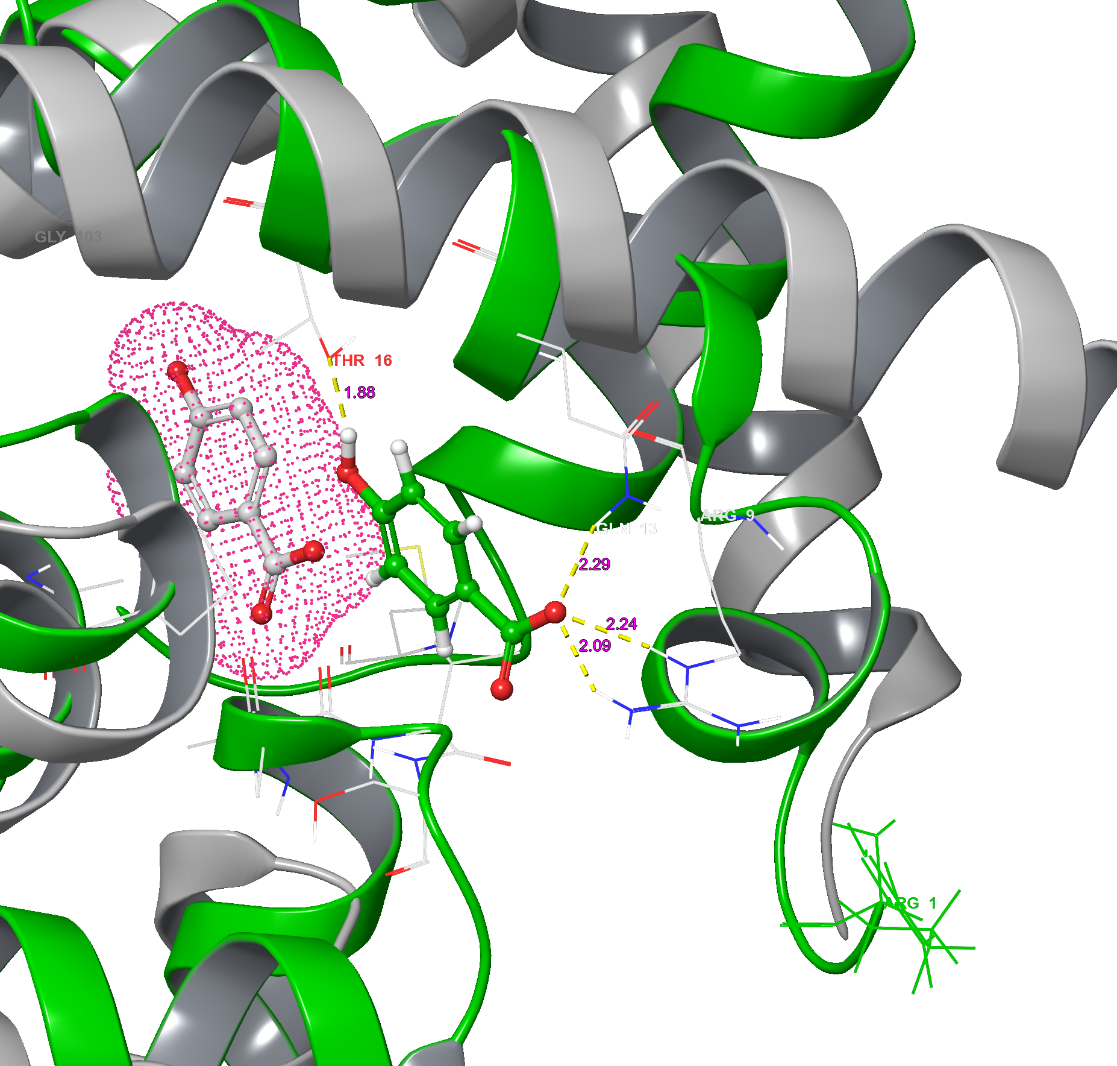

**Figure 2A** The N-terminus of HosA-p-hydroxybenzoic acid complex is shown in green at 98.90ns, superimposed on the grey 0ns reference frame. Yellow dash lines indicate hydrogen bonds, with salmon pink representing the bond distance. The parent p-hydroxybenzoic acid is depicted in pink mesh at the 0ns frame.

**Figure 2B** The N-terminus of HosA-p-hydroxybenzoic acid complex is shown in green at 1.20ns, superimposed on the grey 0ns reference frame. The parent p-hydroxybenzoic acid is depicted in pink mesh at the 0ns frame.

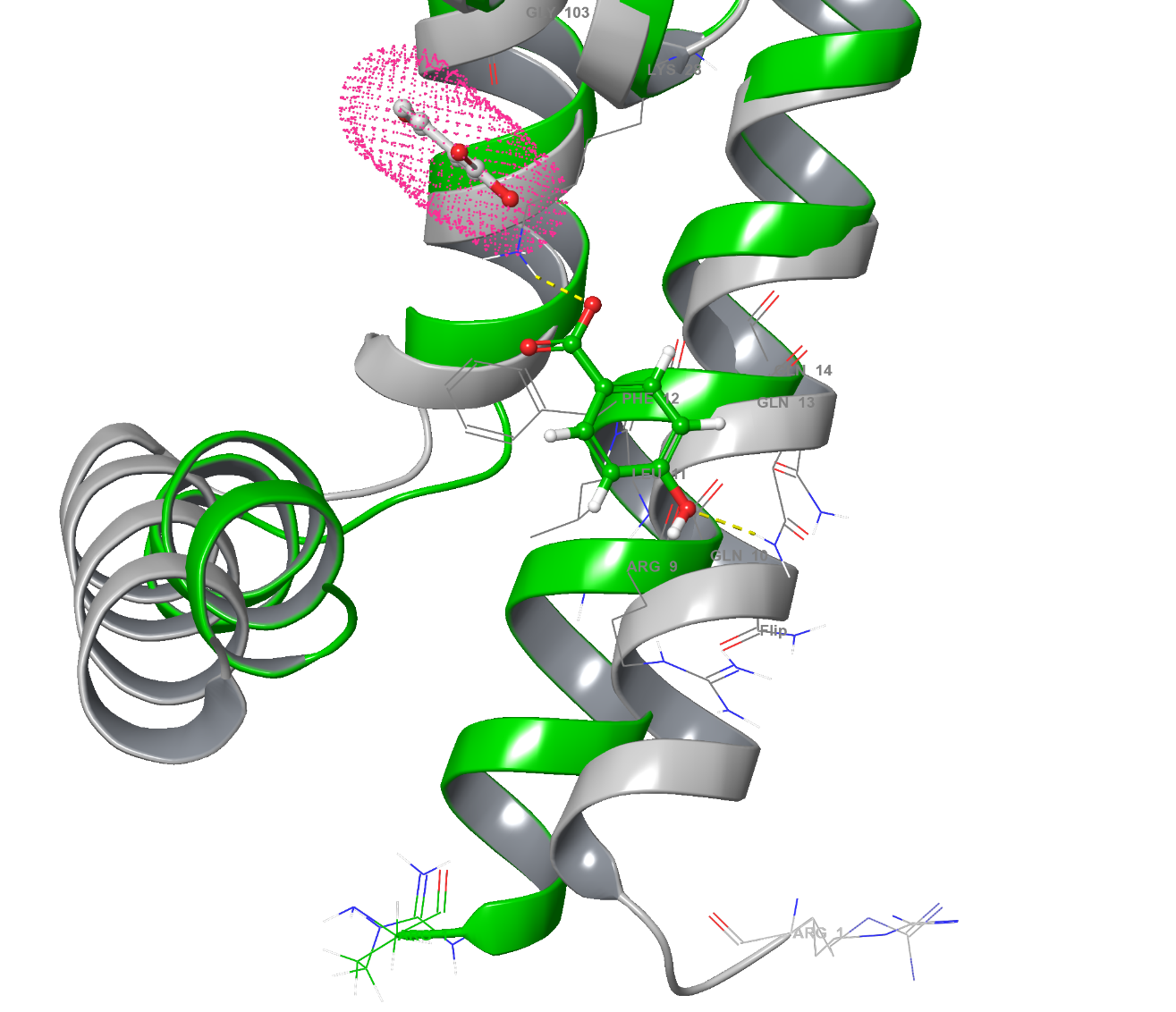

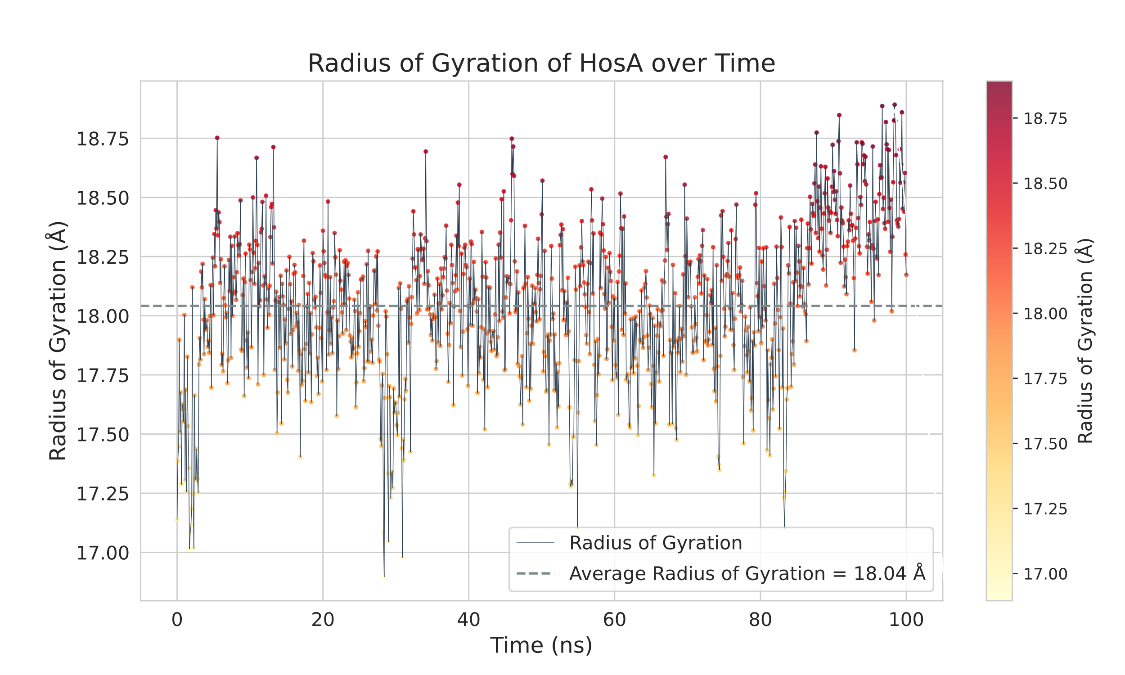

**Figure 3A** The line graph depicts the radius of gyration (Rg) of HosA-apo, with an average value indicated by a dashed grey line.

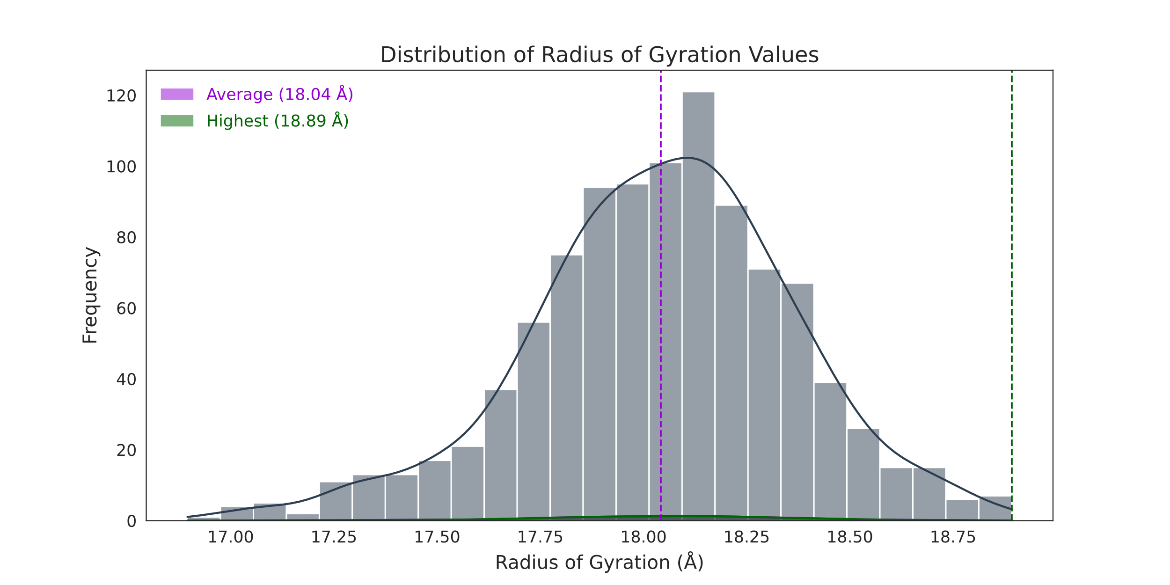

**Figure 3A** The histogram illustrates the distribution of the maximum radius of gyration (Rg) of HosA-apo, represented by a pink salmon color. The average Rg value is denoted by a dashed green line.

**Figure 3A** The histogram illustrates the distribution of the maximum radius of gyration (Rg) of HosA-apo, represented by a pink salmon color. The average Rg value is denoted by a dashed green line.

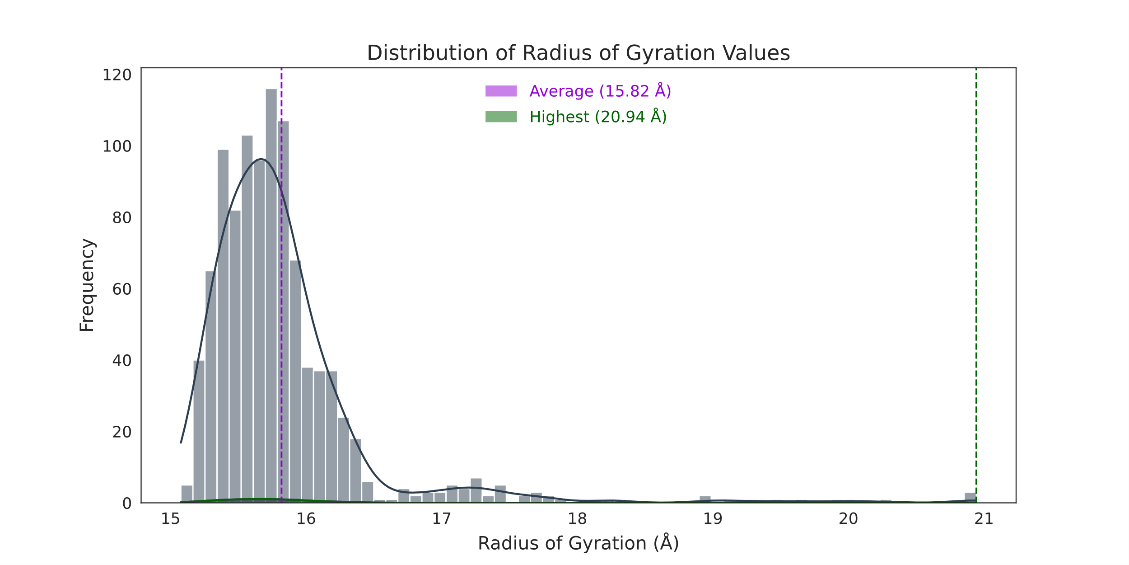

**Figure 3A** The line graph depicts the radius of gyration (Rg) of HosA-apo, with an average value indicated by a dashed grey line.

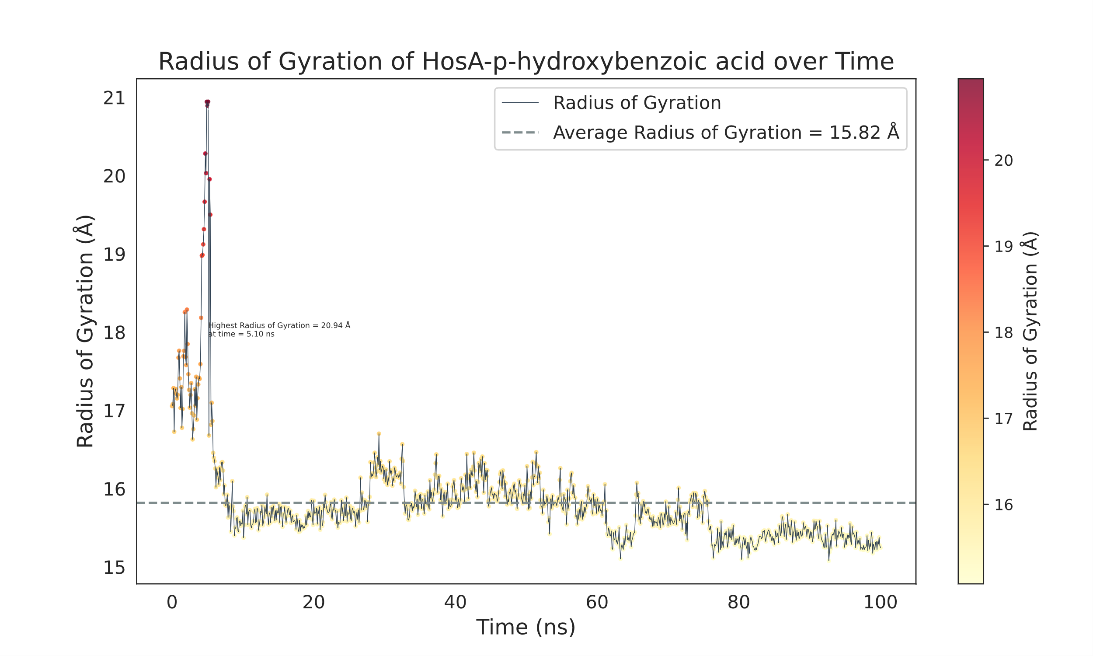

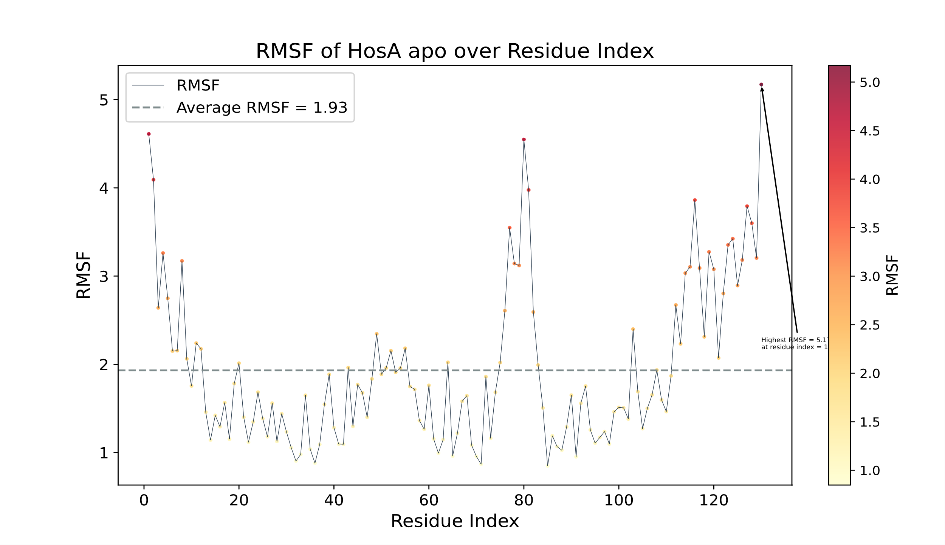

**Figure 4A** The line graph depicts the root mean square fluctuation (rmsf) of HosA-apo, with an average value indicated by a dashed grey line.

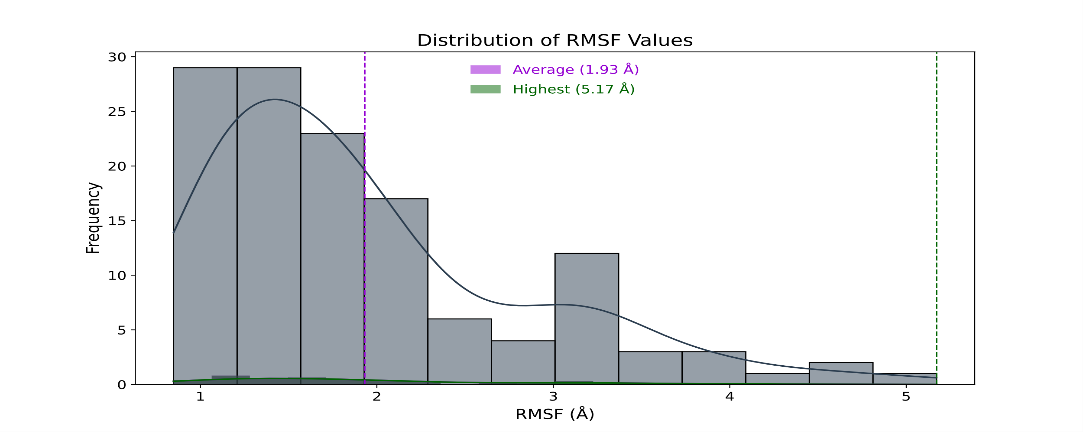

**Figure 4B** The histogram illustrates the distribution of the maximum root mean square fluctuation (rmsf) of HosA-apo, represented by a pink salmon color. The average is denoted by a dashed green line.

**Figure 4C** The line graph depicts the root mean square fluctuation (rmsf) of HosA-complex, with an average value indicated by a dashed grey line.

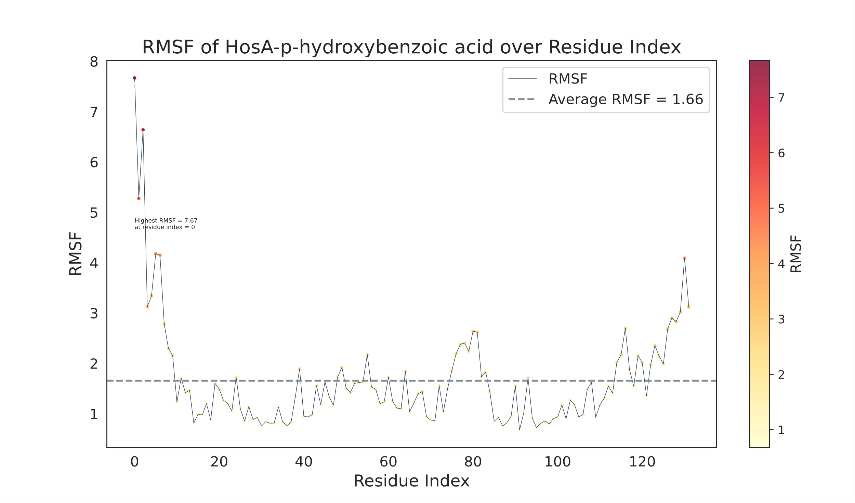

**Figure 4D** The histogram illustrates the distribution of the maximum root mean square fluctuation (rmsf) of HosA-complex, represented by a pink salmon color. The average is denoted by a dashed green line.

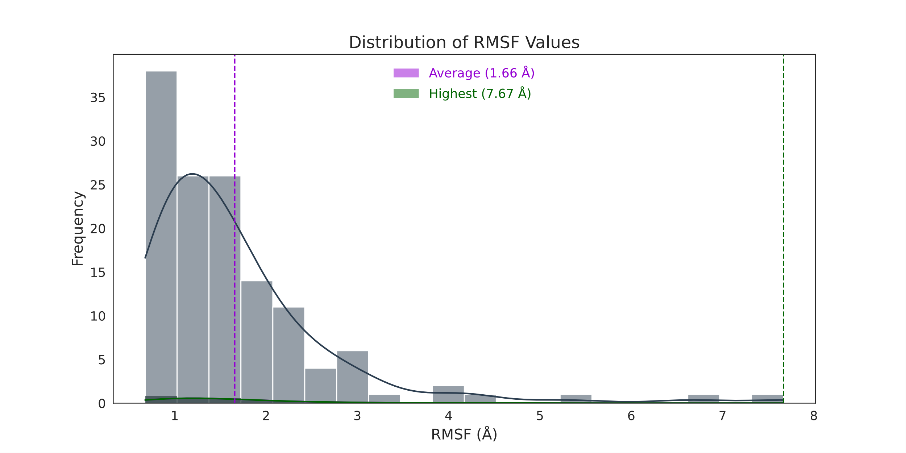

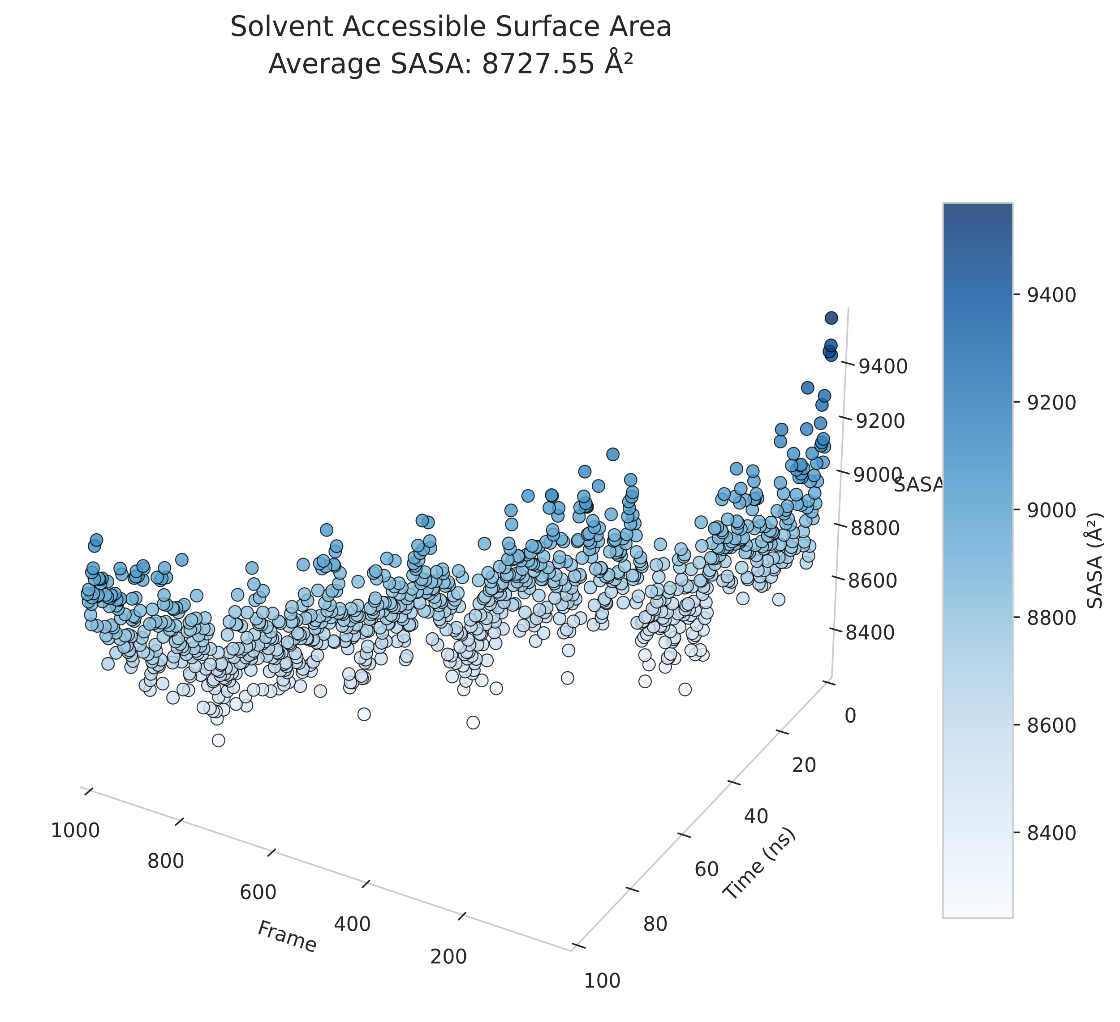

**Figure 5A** The line graph illustrates the variations in solvent accessible surface area (SASA) of HosA-apo. The low SASA values are represented by light grey color, while the high SASA values are depicted by dark teal to blue color.

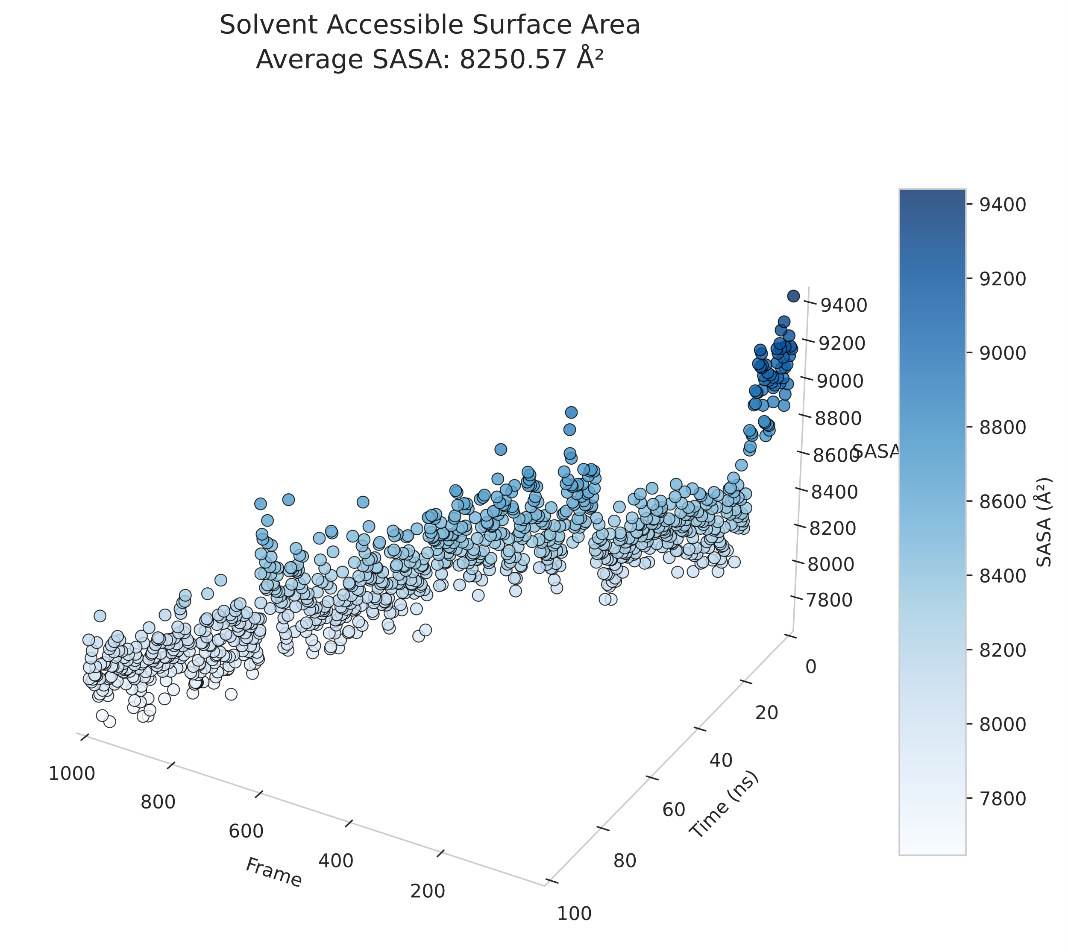

**Figure 5B** The line graph illustrates the variation in solvent accessible surface area (SASA) of the HosA-complex. The areas with low SASA are represented by light grey color, while regions with high SASA are depicted by shades of dark teal to blue.

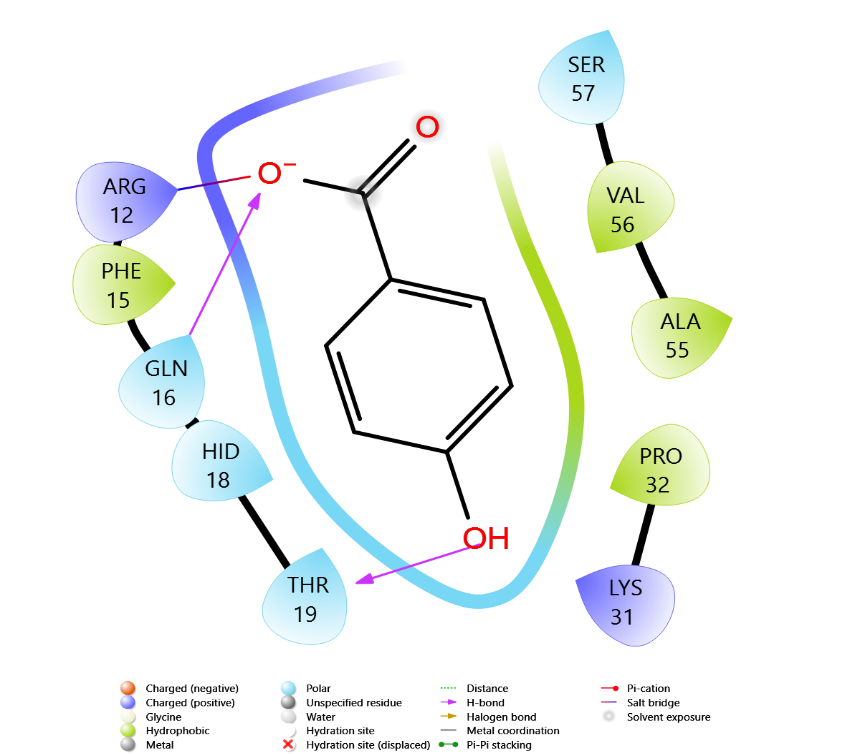

**Figure 6B** Depicts the two-dimensional visualization of the HosA-p-hydroxy benzoic acid complex, highlighting the hydrogen bonds with the highest frequency with ARG12, GLN16 and THR19 observed at the 100 nanosecond time point. The pink lines in the figure indicate the specific hydrogen bond interactions.

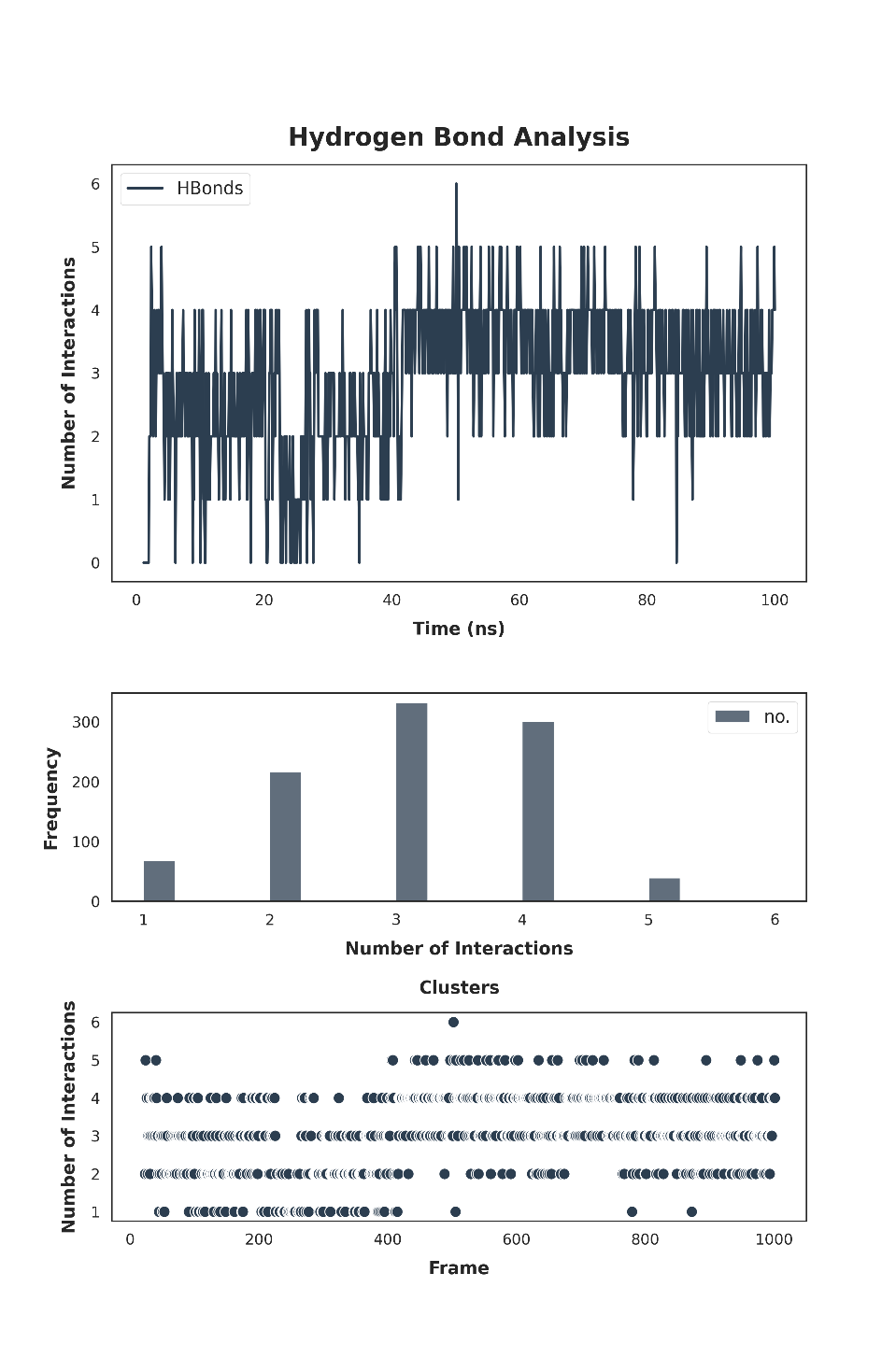

**Figure 6A** Shows a line graph that shows the specific hydrogen bonds formed in a 100ns time frame for HosA-Complex. The graph also highlights the frequency and clustering of these interactions in a visually distinguishable dark teal color.

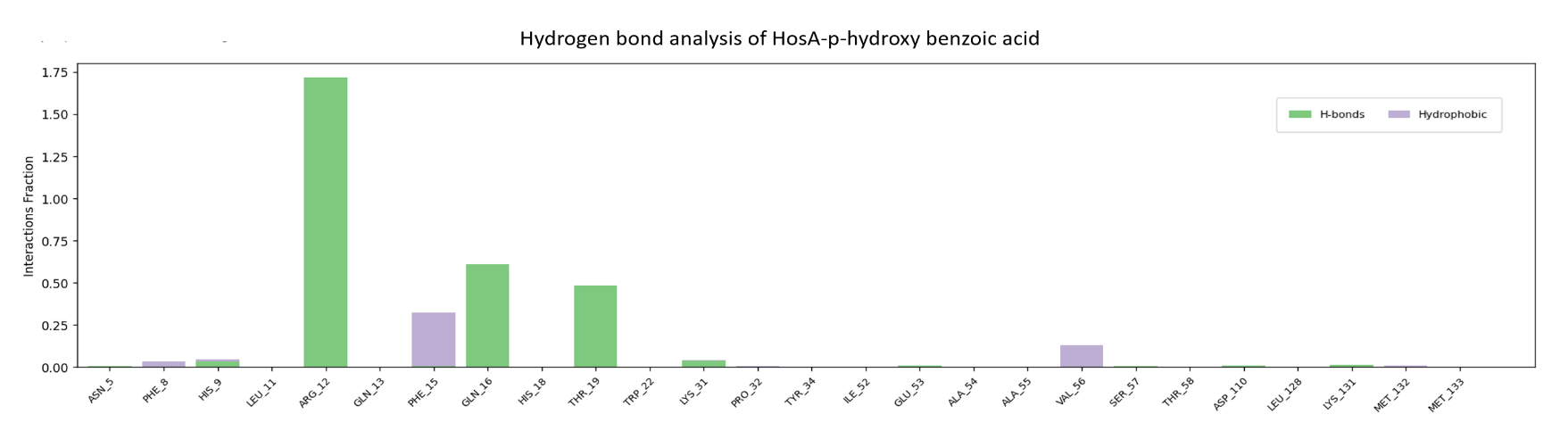

**Figure 6C** Illustrates a bar graph displaying the number of hydrogen bonds formed with high frequency at 100ns among ARG12, GLN16, and THR19. The green bar represents the hydrogen bonds between HosA-p-hydroxy benzoic acid.

**Figure 6D** Illustrates a scatter plot that demonstrates the count of internal hydrogen bonds established in HosA-apo after 100ns,

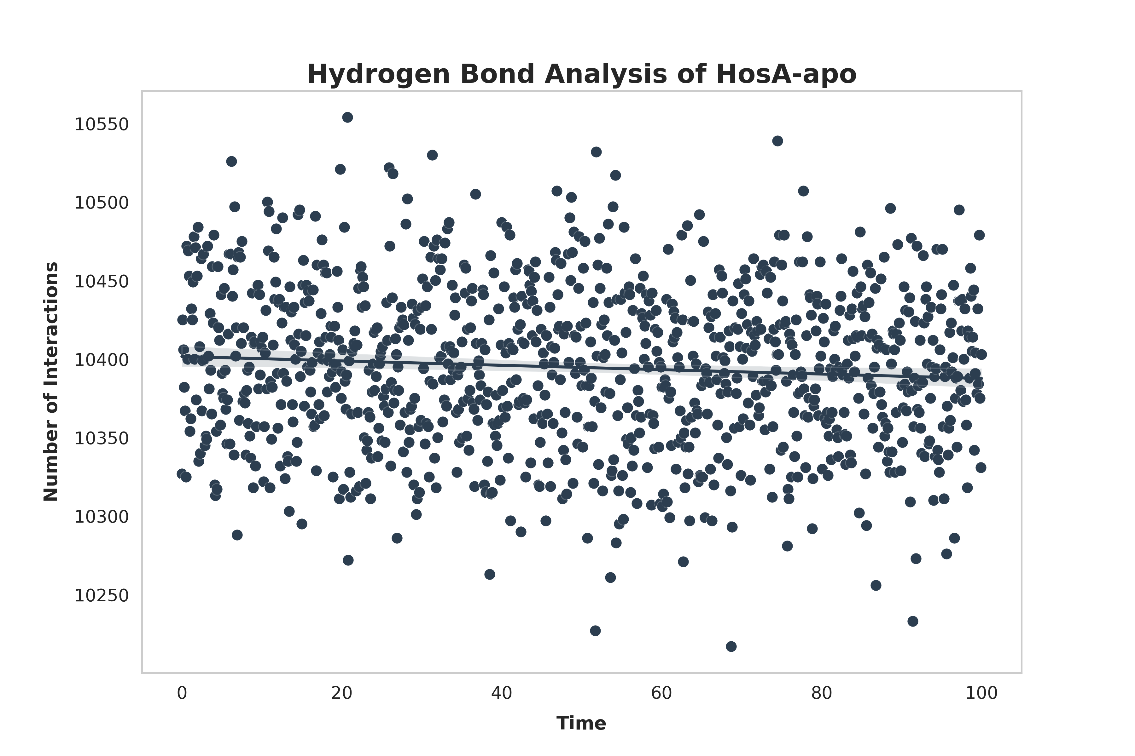

**Simulation Quality Analysis of HosA-Apo vs Complex**

The quality analysis of the molecular dynamics simulation of the HosA-apo and p-hydroxybenzoic acid bound complex was conducted, assessing various descriptive parameters such as total energy, potential energy, volume, pressure, and temperature of the entire system. The results, depicted in Table 1 and Figure 7A-B, were indicative of the structural stability and intermolecular interactions of the protein-ligand complex. The total energy of the HosA-apo structure was found to be -81162.275 (kcal/mol), while that of the HosA-p-hydroxybenzoic acid complex was -67523.161 (kcal/mol). This suggests that the apo structure is more stable than the complex, which is supported by the lower root mean square deviation (rmsd) values and reduced fluctuations observed in the apo structure in Figure 1A-B. Additionally, the potential energy of the complex was lower (-82615.578 kcal/mol) than that of the apo structure (-99128.252 kcal/mol), indicating stronger interactions between the HosA and p-hydroxybenzoic acid. This is further supported by the hydrogen bond analysis depicted in Figure...

Furthermore, the volume of the HosA-apo structure was measured to be 298288.131 Å³, whereas the HosA-p-hydroxybenzoic acid complex had a volume of 249388.461 Å³, showing a significant decrease in volume upon binding of the p-hydroxybenzoic acid. Additionally, the pressure was found to be slightly higher for the complex (1.113 bar) than the apo structure (1.028 bar), suggesting a more compressed system in the presence of the p-hydroxybenzoic acid. Nonetheless, the temperature of both structures was consistent, with a value of 298.705 K, indicating proper equilibration of the system.

**Table 1.** Simulation quality analysis of both (HosA-apo and p-hydroxybenzoic acid complex) full systems at 100 ns. Slope (ps^-1^) was 0 in all cases.

| Descriptive parameters | Average (Apo and Complex) | | Std. Dev. | |
| --- | --- | --- | --- | --- |
| Total energy (kcal/mol) | -81162.275 | -67523.161 | 73.509 | 67.172 |
| Potential energy (kcal/mol) | -99128.252 | -82615.578 | 61.967 | 58.013 |
| Temperature (K) | 298.705 | 298.705 | 0.667 | 0.706 |
| Pressure (bar) | 1.028 | 1.113 | 52.530 | 60.275 |
| Volume (Å³) | 298288.131 | 249388.461 | 319.479 | 297.074 |

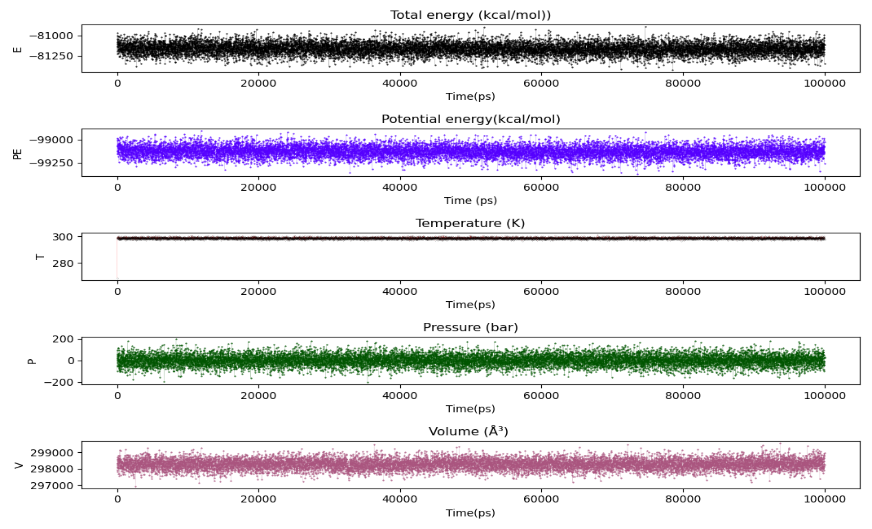

**Figure 7A** Descriptive parameters of the HosA-apo system at 100 ns. E=Total energy(kcal/mol) in black, PE=Potential energy(kcal/mol) in blue, T=Temperature(K) in black, P=Pressure(bar) in dark green and V=Volume (Å³) in salmon color representation

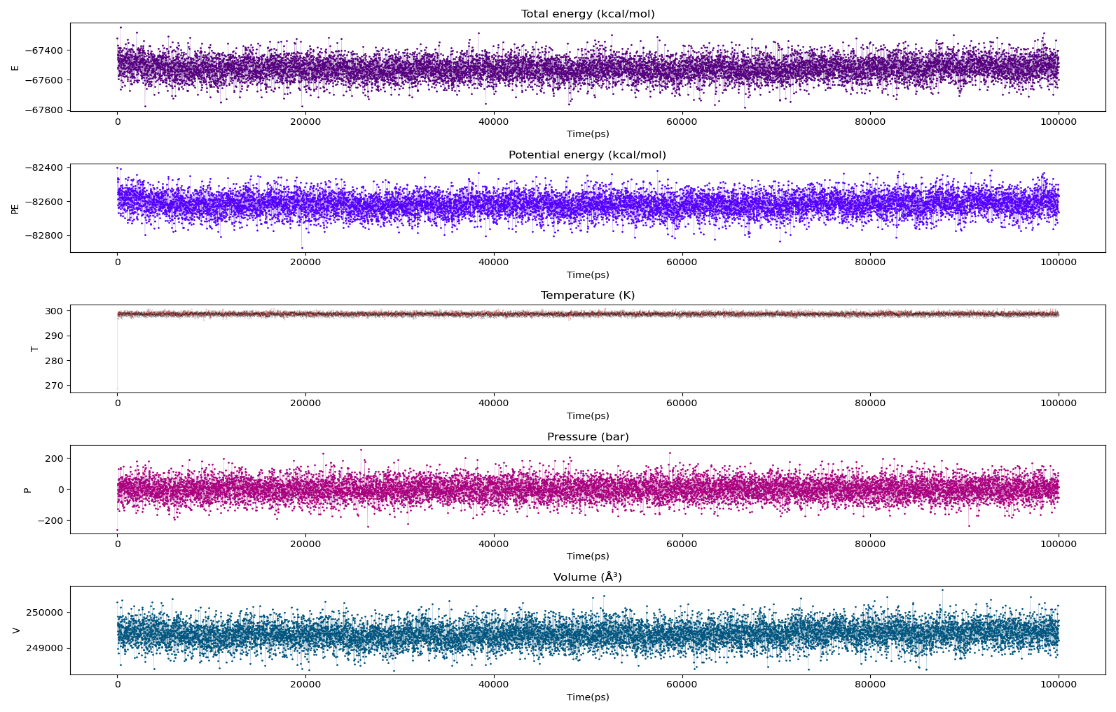

**Figure 7B** Descriptive parameters of the HosA-p-hydroxybenzoic acid complex system at 100 ns. E=Total energy(kcal/mol) in purple, PE=Potential energy(kcal/mol) in blue, T=Temperature(K) in black-red, P=Pressure(bar) in pink and V=Volume (Å³) in teal color representation
