## Supplementary Table 1 for "Horizontally acquired HosA transcription factor bound with 4-hydroxy-benzoic acid exhibits unique tug-of-water dynamics"

### Table 1 *–* Data collection, processing, refinement statistics and model quality parameters for structure of HosA (PHB bound conformation II).

| *Structure of HosA (PHB bound conformation II)* - PDB entry 8YCV | | |
| --- | --- | --- |
| Data Collection | | |
| Synchrotron Facility | INDUS synchrotron (RRCAT, Indore, India) | |
| Beamline | PX-BL21 (Indus-2) | |
| Wavelength (Å) | 0.9789 | |
| **Data Processing** | | |
|  | ***autoPROC/STARANISO*** | ***autoPROC/AIMLESS*** |
| Resolution range (Å)^a^ | 67.17 - 2.16  (2.33 – 2.16) | 54.87 – 2.32  (2.36 – 2.32) |
| Space group | *P* 4_3_ 2 2 | |
| Unit cell parameters  *a, b, c* (Å)  *α, β, ϒ* (º) | 67.17, 67.17, 95.10  90.0, 90.0, 90.0 | |
| Total no. of reflections | 60 800 (1 821) | 63 016 (2 394) |
| No. of unique reflections | 8 907 (445) | 9 954 (461) |
| Multiplicity | 6.8 (4.1) | 6.3 (5.2) |
| Completeness (%) |  |  |
| Spherical | 73.2 (19.3) | 99.6 (99.8) |
| Ellipsoidal | 94.1 (82.7) | - |
| Mean I/σ(I) | 9.4 (1.4) | 8.4 (1.0) |
| *R*_merge_ (%) ^b^ | 13.5 (102.2) | 14.0 (160.6) |
| *R*_meas_ (%) ^c^ | 14.7 (117.2) | 15.2 (178.2) |
| *R*_pim_ (%) ^d^ | 5.5 (55.3) | 5.9 (75.3) |
| *CC*_1/2_ (%) ^e^ | 99.7 (46.0) | 99.6 (44.8) |
| **Refinement** | | |
|  | ***autoPROC/STARANISO*** | ***autoPROC/AIMLESS*** |
| *R*_work_ (%) ^f^ | 20.18 (29.11) | - |
| *R*_free_ (%) ^g^ | 23.19 (32.18) | - |
| RMSD Bonds (Å) ^h^ | 0.003 | - |
| RMSD Angles (º) ^h^ | 0.499 | - |
| Number of atoms |  | - |
| Protein residues | Ala2 – Asn134 | - |
| Non-hydrogen atoms | 1 146 | - |
| Macromolecules | 1 071 | - |
| Ligands | 24 | - |
| Waters | 51 | - |
| Ramachandran plot |  |  |
| Most favoured (%) | 99.24 | - |
| Outliers (%) | 0.00 | - |
| Rotamer outliers (%) | 0.00 | - |
| Clashscore ^i^ | 0.45 | - |
| Molprobity score ^j^ | 0.66 | - |
| Average *B*-factors (Å^2^) | 41.99 |  |
| Protein | 42.18 | - |
| Ligands | 41.80 | - |
| Solvent | 38.03 | - |

^a^ Information in parenthesis refers to the last resolution shell.

^b^ $R_{merge}=\sum_{hkl} \sum_{i} |I_{i}\left( hkl \right)-\bar{I\left( hkl \right)|}/\sum_{hkl} \sum_{i} I_{i}\left( hkl \right)$.

^c^$R_{meas}= \sum_{hkl} [N/{\left( N-1 \right)]}^{\frac{1}{2}} \sum_{i} |I_{i}\left( hkl \right)-\bar{I\left( hkl \right)}| /\sum_{hkl} \sum_{i} I_{i}\left( hkl \right).$

^d^ $R_{p.i.m}= \sum_{hkl} [1/{\left( N-1 \right)]}^{\frac{1}{2}} \sum_{i} |I_{i}\left( hkl \right)-\bar{I\left( hkl \right)}| /\sum_{hkl} \sum_{i} I_{i}\left( hkl \right)$.

^e^ *CC*_1/2_ as described in Karplus & Diederichs (2012). Science, 336(6084): 1030–1033.

^f^ $R_{work}= \sum_{h} \sum_{k} \sum_{l} \frac{\left\{ \left| \left| \left. F_{o}(h,k,l) \right| \right. \right.\left. -\left| \left. F_{c}(h,k,l) \right| \right| \right\} \right.}{\sum_{h} \sum_{k} \sum_{l} \left| F_{o}(h,k,l) \right|}$, where F_o_ and F_c_ are the observed and calculated structure factors for reflection h, respectively.

^g^ *R*_free_ was calculated the same way as R_work_ but using only 5% of the reflections which were selected randomly and omitted from refinement.

^h^ RMSD, root mean square deviation.

^i^ Clashscore is the number of unfavourable all-atom steric overlaps ≥ 0.4Å per 1000 atoms*.* Word *et al*. (1999). Mol Biol, 285(4):1711-33.

^j^ *MolProbity* score provides a single number that represents the central *MolProbity* protein quality statistics; it is a log-weighted combination of Clashscore, Ramachandran not favoured and bad sidechain rotamers, giving one number that reflects the crystallographic resolution at which those values would be expected.
